# Early life lipid overload in Native American myopathy is phenocopied by *stac3* knock out in zebrafish

**DOI:** 10.1101/2023.07.26.550753

**Authors:** Rajashekar Donaka, Houfeng Zheng, David Karasik

**Author notes:** Corresponding author: David Karasik, PhD The Musculoskeletal Genetics Laboratory Azrieli Faculty of Medicine, Bar-Ilan University, Safed 1311502, Israel.

## Abstract

Understanding the early stages of human congenital myopathies is critical for proposing strategies for improving skeletal muscle performance by the functional integrity of cytoskeleton. SH3 and cysteine-rich domain 3 (Stac3) is a protein involved in nutrient sensing, and is an essential component of the excitation-contraction (EC) coupling machinery for Ca^2+^ releasing. A mutation in *STAC3* causes debilitating Native American myopathy (NAM) in humans, and loss of this gene in mice and zebrafish resulted in death in early life. Previously, NAM patients demonstrated increased lipids in skeletal muscle biopsy. However, elevated neutral lipids could alter muscle function in NAM disease via EC coupling apparatus is yet undiscovered in early development.

Here, using a CRISPR/Cas9 induced *stac3* knockout (KO) zebrafish model, we determined that loss of *stac3* led to muscle weakness, as evidenced by delayed larval hatching. We observed decreased whole-body Ca^2+^ level at 5 days post-fertilization (dpf) and defects in the skeletal muscle cytoskeleton, i.e., F-actin and slow muscle fibers at 5 and 7 dpf. Homozygous larvae exhibited elevated neutral lipid levels at 5 dpf, which persisted beyond 7 dpf. Myogenesis regulators such as *myoD* and *myf5*, were significantly altered in *stac3^-/-^* larvae at 5 dpf, thus a progressive death of the KO larva by 11 dpf.

In summary, the presented findings suggest that *stac3^-/-^* can serve as a non-mammalian model to identify lipid-lowering molecules for refining muscle function in NAM patients.

## Introduction

The musculoskeletal system provides a unique framework for bone, skeletal muscle, and connective tissue for enabling mobility and withstanding mechanical load generated in daily activities of human life. Skeletal muscle (∼40% in the human body mass) serves to provide anatomical support for the bones/internal organs, to produce contractions and to store energy for locomotion, all essential structures and processes for the survival of an organism^1^. Defects in the genetic makeup or metabolic activity of skeletal muscle result in dystrophy, myopathy, cachexia and/or sarcopenia. Among these muscle diseases, myopathies have often been reported as heterogeneous entities that include channelopathies, inflammatory, metabolic, mitochondrial, and myotoxic diseases, which affect 1.62 in 100,000 individuals of all ages across the globe^2, 3, 4, 5^.

Muscle weakness is one of the key hallmarks of congenital myopathies (1 in 26,000)^6^, while its further consequences are escalated fibrosis, fat infiltration, and degenerating myofibers^7^. The cytoskeleton of muscle tissue is composed of filamentous actin (F-actin) and myosin. Of late, it was shown that the structural position of fast muscle cell fusion was guided by slow muscle fibers in myotome formation^8^. In vertebrates, slow muscle fibers originate from somitic mesoderm as one of the embryonic lineages; these embryonic lineages further differentiate into the first muscle fiber types to perform longer contractions by firing action potential^8, 9, 10^. In early life, human myopathies are usually associated with accumulation of disorganized actin, α-actinin and myosin, and failure of myofibrillar assembly in fast-twitch muscle fibers^11^.

Historically, Native American myopathy (NAM) was identified in the Lumbee Indian infant population of North Carolina, USA. The genetic predisposition to NAM was determined to be caused by homozygosity for a variant G>C in exon 10 of *STAC3*, a gene highly expressed in skeletal muscle. The STAC3 protein contains a Src homology 3 (SH3) and cysteine-rich C1 domains. Clinically, other genetic mutations in *STAC3* result in congenital anomalies, such as cleft palate, micrognathia, *talipes equinus* (club foot), and arthrogryposis also found in NAM patients. Besides these skeletal abnormalities, NAM patients demonstrate increased levels of neutral lipids in muscle biopsy and it was reported that one-third of patients (36%) of disease afflicted individuals die before the age of 18 years^12, 13^. Of late, it was suggested that C1 domain of STAC3 is likely to bind the lipids^14^. Multiple studies have focused on investigating the regulation of *STAC3* in NAM. Among all, *Stac3* knockout mice had a higher percentage of type I (oxidative) muscle fibers and exhibited poor musculoskeletal performance^15^. Transcriptional regulation of *Stac3* suggests that nutrient balance is a prerequisite for cytoskeletal integrity and myogenesis via energy metabolism^16^. However, the genetic etiology and pathophysiology of slow skeletal muscle weakness in NAM remain largely elusive.

Conventionally, lipids play a significant role in energy homeostasis, cellular structure and cellular signaling, while lipid composition is crucial for ion channel activity and maintenance of membrane receptor conformation^17, 18^. Oftentimes, alterations in lipid metabolism account for metabolic muscle diseases^19^. Importantly, storage of complex lipid molecules or glycogen content in skeletal muscle can modify the calcium (Ca^2+^) release at excitation-contraction (EC) coupling apparatus on sarcoplasmic reticulum (SR). Interestingly, calcium-sensing dihydropyridine receptor (DHPR) and calcium-releasing ryanodine receptor (RYR1) are under the control of STAC3 for intracellular functions including muscle contractions^20, 21^. These physiological actions involving the EC apparatus underscore the absolute necessity of STAC3 for Ca^2+^ and nutrients homeostasis.

One decade ago, the first *stac3* knockout study in zebrafish revealed that a mutation in the *stac3* gene of zebrafish (ZF) larvae impeded muscle function through EC coupling-driven defects in the motor system in early life. STAC3 regulates the amount and stability of DHPR and/or RYR1 at skeletal muscle triads by protein trafficking^22^. Markedly, the nutrient load (neutral lipids) accelerates degeneration of filamentous actin (F-actin) and slow muscle myosin (Smyhc1) fibers, which exemplify harmful behaviour of lipids in the maturation of skeletal muscle fibers. Thus, identification of genetically-driven metabolic peculiarities, i.e., in lipid composition, may be vital for preventing the loss of muscle fibers. Such disease-associated fatty acids may serve as biomarkers for the early diagnosis of the disease or primary clinical indicators of treatment success of NAM patients.

Of late, human myopathies have been modeled by knocking-out disease-causing genes using CRISPR/Cas9 technology. In parallel, ZF serve as remarkable teleost animal model for skeletal muscle research^23^ and as a non-mammalian model for myopathies. Inherently, ZF acquired multiple advantages, including high fecundity, external fertilization, and quick transformation of one-to-multiple somite stages of the larva, which could provide substantial clinical and histopathological information on the muscle disease^24, 25^. The current study generated *stac3^-/-^* zebrafish using CRISPR/Cas9 technology. Remarkably, the *stac3^-/-^* fish resemble human phenotypes of Native American myopathy with increased neutral lipids level in the larvae. This metabolic disbalance was further evident in structural and functional defects in zebrafish, which seemingly drive failure of *stac3^-/-^* larvae’s musculature and their early demise.

## Methods

### Animal husbandry

Danio rerio zebrafish strains were maintained at 28 °C, under a 14:10 light: dark cycle and their offspring was propagated until 6 dpf in system water containing 0.1% methylene blue (embryo water) and then transferred to the system (nursery) or kept in the Petri dishes sans methylene blue, for beginning of feeding. All animal experiments were conducted in accordance with the Faculty of Medicine’s Zebrafish Facility, Bar-Ilan University, Israel, Institutional Animal Care and Ethical Committee (IACUC) guidelines. The protocol was approved by the committee (protocol #53-08-2020).

### Generation of *stac3*^-/-^ by CRISPR/Cas9 system

The strategy for establishing a stable zebrafish knockout followed a published CRISPR/Cas9 protocol^26^. In brief, the guide RNA (gRNA) was designed to target sequence 5’-AGTTCTGTGACGTCTGCGCACGG-3’ in exon 4 of the *stac3* gene. The *FspI* restriction enzyme (New England BioLabs, Cat#R0135S) was utilized to distinguish between mutant, wild type and heterozygous fish (Supplementary materials and methods). At the one-cell stage (30-45 minutes post-fertilization), wild type zebrafish embryos of AB strain (F0) were injected with a mixture of gRNA, crRNA, tracer RNA and Cas9 protein (Abcam, Cat#224892), at total amount of 300 µg/µL, using a pneumatic Pico Pump (WPI, Worcester, MA, USA), and embryos were transferred into the incubator.

### Genotyping and sequencing

After 24 hours, DNA was extracted from of subset of pooled gRNA mix-injected embryos. A polymerase chain reaction (PCR) was carried with forward primer: 5’-GTGTTTTCAACGTTAGTTCTGCTG-3’ and reverse primer: 5’-TGGCAAGAACAGTTTACCTTAACA-3’. The reactions were performed using two programs: (A) DNA was denatured at 94 °C for 3 seconds, annealed at 63 °C for 3 seconds and extended at 72°C for 5 cycles, 3 seconds each. (B) DNA was denatured at 94 °C for 3 seconds, annealed at 60 °C for 3 seconds and extended at 72 °C for 37 cycles, 3 seconds each, followed by final extension at 72 °C, for 3 minutes. Products of PCR amplification and restriction digestion were examined on a 2% agarose gel. The 10-base-pair (bp) deletion was verified by DNA sequencing (Hy-lab, Rehovot, Israel and Macrogen, Amsterdam, Netherlands).

For generation of a stable knockout, CRISPR/Cas9-targeted larvae were grown in a 1-liter (L) tank for 35 days and then maintained in 3-L tank in a system water. At 2 months post-fertilization (mpf), fish were genotyped to identify a founder that carried a mutation. Germline transmission of a mutation in F1-progeny was deciphered by crossing the adult founders (F0) to WT. To acquirethe 10 bp deletion in F2-progeny, adults (F1) were out-crossed to WT. To minimize an off-target effect, adult fish carrying the 10 bp deletion (F2) were further outcrossed to WT. The *stac3*^+/+^, *stac3*^+/-^, and *stac3*^-/-^ siblings were obtained by intercrossing *stac3*^+/-^ F3 adult parents.

### Calculating hatching percentage of *stac3*^-/-^ zebrafish larvae

Mass spawning of adult *stac3*^+/-^ parents produced all three genotypes: *stac3^+/+^*, *stac3^+/-^* and *stac3^-/-^* according to the Mendelian ratio. After spawning, embryos were collected and grown in a Petri plate containing embryo water. During the course of somite to muscle fiber transition, the number of hatched and unhatched larvae was recorded at 3, 4 and 5 dpf, and delayed hatching of the larvae was documented by bright field light microscope imaging, while their genotypes were verified by DNA sequencing of pooled sample (n=10) for each group. Delayed hatching observations and analyses were conducted on at least three independent clutches of parent’s spawning. We saw skeletal peculiarities early on, thus we identified larvae as “normal” (*stac3* wild type and heterozygous siblings) and “deformed” (*stac3* knockout) for the ensuing study.

As a rule, larvae were examined under the light microscope every 24 hours. The total number of embryos produced by *stac3*^+/-^ parents was recorded. Dead larvae at (4-11 dpf) were extracted from Petri plates and genotyped. In all experiments, larvae were euthanized in 0.4% methane sulfonate (Sigma-Aldrich, MS-222) with subsequent immersion in ice-cold water for 15 minutes.

### Whole-body Ca^2+^ measurement during larval development

Total body calcium level was determined for zebrafish larvae groups, i.e. *stac3^+/+^* and *stac3^+/-^*: n=90 and *stac3^-/-^*: n=60). After euthanasia, larvae were washed gently with deionized water and dried for 1 hour at 65 °C. Then, samples were digested with 80-90 µL 1M Tris-Cl (pH 8.0), at 95 °C, overnight. Digested samples were centrifuged at 13,000 rpm for 5 minutes and the supernatant was collected in 1.5 mL Eppendorf tube. Calcium amount was detected using the Colorimetric Calcium Assay Kit (Abcam, Cat#102505), according to the manufacturer’s instructions. Optical density of each sample was measured at 575 nm with absorption spectrometry (Tecan-Plate Reader-Spark Cyto, DKSH technology, Bangkok, Thailand).

### RNA extraction

We extracted RNA from *stac3* wild type and heterozygous siblings and *stac3* knockout for gene expression analysis at 4 and 5 dpf. After euthanasia, pooled (n=10) larvae were homogenized with a pestle and motor mixer (Thomas Scientific, Swedesboro, NJ, USA, Cat#47747-370) in 350 µL Trizol (Sigma-Aldrich). RNA was extracted in biological triplicates, using the Direct-Zol RNA kit (Zymo Research, Tustin, CA, USA).

### Real-time quantitative PCR (RT-qPCR)

cDNA (500-1000 ng) was synthesized using the Takara PrimeScript Kit (Takara, Mountain View, CA, USA). Gene expression was quantified using the PowerUp SYBR Green Master Mix and a ViiA™7 Dx qPCR Instrument (Thermo Fisher Scientific, Waltham, MA, USA, Cat#4453534). The target gene expression (TableS1) was normalized to the endogenous control *rpl32* (ribosomal protein l32). For each expression analysis, a non-template control (NTC) was included along with technical triplicates. Delta-delta-threshold cycle (DDCT) values were plotted on the graph as relative gene expression.

### Whole-mount staining of zebrafish larvae

At 5 and 7 dpf, *stac3^+/+^* and *stac3^+/-^*: n=15 and *stac3^-/-^*: n=15 larvae were fixed in 4% paraformaldehyde (PFA, Sigma-Aldrich) either for 1 hour at RT or overnight at 4 °C. Staining was performed using a published protocol^27^. Slow muscle fibers were stained with the anti-Myhc (F59, Santa Cruz Biotechnology, Dallas, TX, USA, Cat#sc-32732) antibody (1:1000 dilution). F-actin fibers were stained with Phalloidin (iFluor 488, Cat#Ab176753) (1:1000 dilution), followed by secondary antibody (Sigma-Aldrich).

### Neutral lipid visualization by Oil Red O (ORO) staining

ORO (Sigma Aldrich, Cat#0-0625) powder 0.05 g was dissolved in 10 mL isopropanol and then incubated for 10 minutes at RT. The ORO working solution was prepared by adding three parts of filtered ORO stock to the two parts of distilled water. Muscle section-containing slides were air-dried for 2-3 hours at RT, washed with 1x PBS for 2 minutes and air-dried again for 2-3 hours. The slides were incubated in 500 µL ORO working solution for 30 seconds, then washed with tap water and air-dried for another 3 hours at RT, before mounting them in 100% glycerol.

### Muscle histology

#### Embedding of zebrafish larva

At 5 dpf, larvae were fixed in 4% PFA for 3-6 days, at 4 °C, and washed 3x with PBS for 15 minutes. Then, they were soaked in 10% sucrose for 48 hours at RT, and then embedded with Tissue Tek (Sakura, Torrance, CA, USA) in 12×12 cm embedding molds (PEEL-A-WAY; Cat#70181). The embedding molds were frozen in liquid nitrogen for about 2-3 hours, and then stored in aluminum foil, at −20 °C. Muscle sections were cut serially to 20 μM thickness with a cryostat (Leica CM1950), and sections were collected on 26X76 mm/1-1.2 mm adhesion microscope slides (BAR-NAOR Ltd., Tel Aviv, Israel, Cat#BN93080C).

#### Hematoxylin and eosin staining

Paraffin-based horizontal muscle sections were dehydrated 3x in xylene for 2 minutes, and then dehydrated in 100% alcohol, before being washed with 80% alcohol and tap water. Slides were then stained with both hematoxylin and eosin solution using the LEICA AUTOSTAINER XL at the Department of the Pathology, Ziv Hospital, Safed, Israel.

#### Locomotion of *stac3*^-/-^ larvae

To quantify locomotion function of *stac3* KO, *stac3^+/+^* and *stac3^+/-^* (n=48) and *stac3^-/-^* (n=22) larvae were grouped at 5 dpf. Tracking was performed with a DanioVision system (Noldus Information technology, Wageningen, NL). For habituation, larvae were individually placed before the test, in 96-well-plates containing 200 µL embryo water (A1-D12: *stac3^+/+^* and *stac3^+/-^* and E1-F10: *stac3^-/-^*) and incubated for 30 minutes at 28 °C. The locomotion tracking strategy included 15 minutes light on, 5 minutes light off, 5 minutes light on. Total distance moved by larvae was averaged for every one-minute represented by a dot on the graph under each (light/dark) condition. After the test, larvae genotype was confirmed using the *FSPI* enzyme digestion.

#### Senescence-associated (SA) β-galactosidase staining

To assess the effect of senescence on early life muscle fiber growth and development in zebrafish, larvae at 6 dpf were classified as normal or deformed and fixed in 4% PFA, at 4 °C, overnight. Larvae were washed with phosphate buffered saline (PBS pH 7.4, followed by PBS pH 6.0), 3x for 10 minutes each. Next, larvae were incubated with 1% X-Gal solution [in PBS pH 6.0: 1 mg of 5-bromo-4-chloro-3-indolyl beta-D-galactosidase (X-Gal)] per mL, 5 mM K^3^Fe[CN]^6^, 5 mM K^4^Fe[CN]^6^,150 mM NaCl and 2 mM MgCl^2^ (Sigma-Aldrich) at 37 °C, overnight^28^. Images were acquired with a bright field microscope.

#### Skeletal muscle defects observed by birefringence assay in zebrafish larvae

*stac3^+/+^* and *stac3^+/-^* control siblings and *stac3^-/-^* larvae were anesthetized on 6 dpf and positioned to allow a dorso-ventral view, in 3% methylcellulose. Muscle fiber architecture was assessed, as described in the published protocol^29^ at the Faculty of Marine Sciences, Ruppin Academic Center, Mikhmoret, Israel.

#### Caffeine treatment of zebrafish larvae

At 5 and 7 dpf, *stac3^+/+^* and *stac3^+/-^* control siblings and *stac3^-/-^* larvae (sample sizes from 25 to 30 animals per group), were subjected to caffeine treatment (Sigma-Aldrich, Cat#C0750). Stock caffeine 1M solution was prepared in double distilled water, and a working concentration of 5mM caffeine (v/v) was subsequently diluted in 1L system water. Each petri dish contained either 30mL of working solution of caffeine or embryo water as no-treatment control solution. Survival of the larvae was monitored for 4 days (96 hours), and each day dead larvae were visually recorded by using the light microscope (then discarded).

#### Image processing

Embryo hatching, developmental defects, and whole-mount Oil Red O staining of neutral lipids were imaged using a Leica microscope (M165 FC). Muscle histology imaging was performed with an automated upright slide-scanning microscope (Axio Scan.Z1, Carl Zeiss Microscopy, Oberkochen, Germany) with a 20×/0.95 objective at z-planes of 0.5mm. Fluorescent images were taken with upright microscope (Apotom.2, Carl Zeiss, Jena, Germany).

#### Data analysis

Most of the experimental data were analyzed with Prism (version 9, GraphPad Software, Inc., La Jolla, CA, USA) and some data sets were processed in Excel. All graphs present means ± standard deviation values. Two independent groups were compared using the student’s t-test. Larvae locomotor activity was analyzed by one-way ANOVA. Survival rate of knockout fish was determined by Prism, using the Log-rank test. In all the analyses, statistical significance was defined as a p<0.05.

## Results

### 1. Generation of CRISPR/Cas9 based *stac3* knockout line in zebrafish to study NAM disease process

Loss of *STAC3* leads to severe skeletal abnormalities in both mice and zebrafish, although a mechanism by which *stac3* mutants die at an early age^22, 15^ is unknown. Further, morbidity in NAM patients had associated with increased lipid levels in skeletal muscle. To study the relation between early muscle fibers development and lipid metabolism, CRISPR/Cas9 system was applied to induce a global knockout of exon 4 of *stac3*. A 10-base-pair deletion in ZF was generated, which led to a frame shift which created a stop codon at V78 and produced a truncated protein (80 aa total length) (Fig1A, B). The targeted knockout in *stac3* gene was confirmed by both *FspI* restriction enzyme digestion and sequencing of genomic DNA (Fig1C and FigS1A, B). Adult *stac3* heterozygous (*stac3^+/-^*) parents were then inter-crossed to obtain homozygous (knockout) *stac3^-/-^* offspring. Further, *stac3^+/-^* progeny were “blindly” characterized as we saw congruence with skeletal phenotypes at the early development of the homozygous larvae. ZF larvae from 2-3 independent clutches were then selected based on their morphology, - either normal (healthy; *stac3^+/+^* and *stac3^+/-^*) or deformed (*stac3^-/-^*) - for functional characterization of skeletal muscle tissue at the early life. All unhatched larvae were genotypically confirmed as *stac3* knockouts (FigS2A-C). At 3 days post-fertilization (dpf), *stac3^-/-^* larvae exhibited multiple congenital skeletal muscle defects, such as bending at head, trunk, and tail regions, and were distinct from *stac3^+/+^* and *stac3^+/-^* control siblings (FigS1C). RT-qPCR analysis showed that the relative gene expression of *stac3* was significantly (p<0.05) downregulated in knockout compared to *stac3^+/+^* and *stac3^+/-^* control siblings at 5 dpf (Fig1D). These findings align with previous observations in *stac3* mutant mice and ZF; we therefore utilized *stac3^-/-^* zebrafish for identifying genetically-driven metabolic regulators, which could be potentially associated with human muscle disease (congenital myopathies) pathology.

**1.**
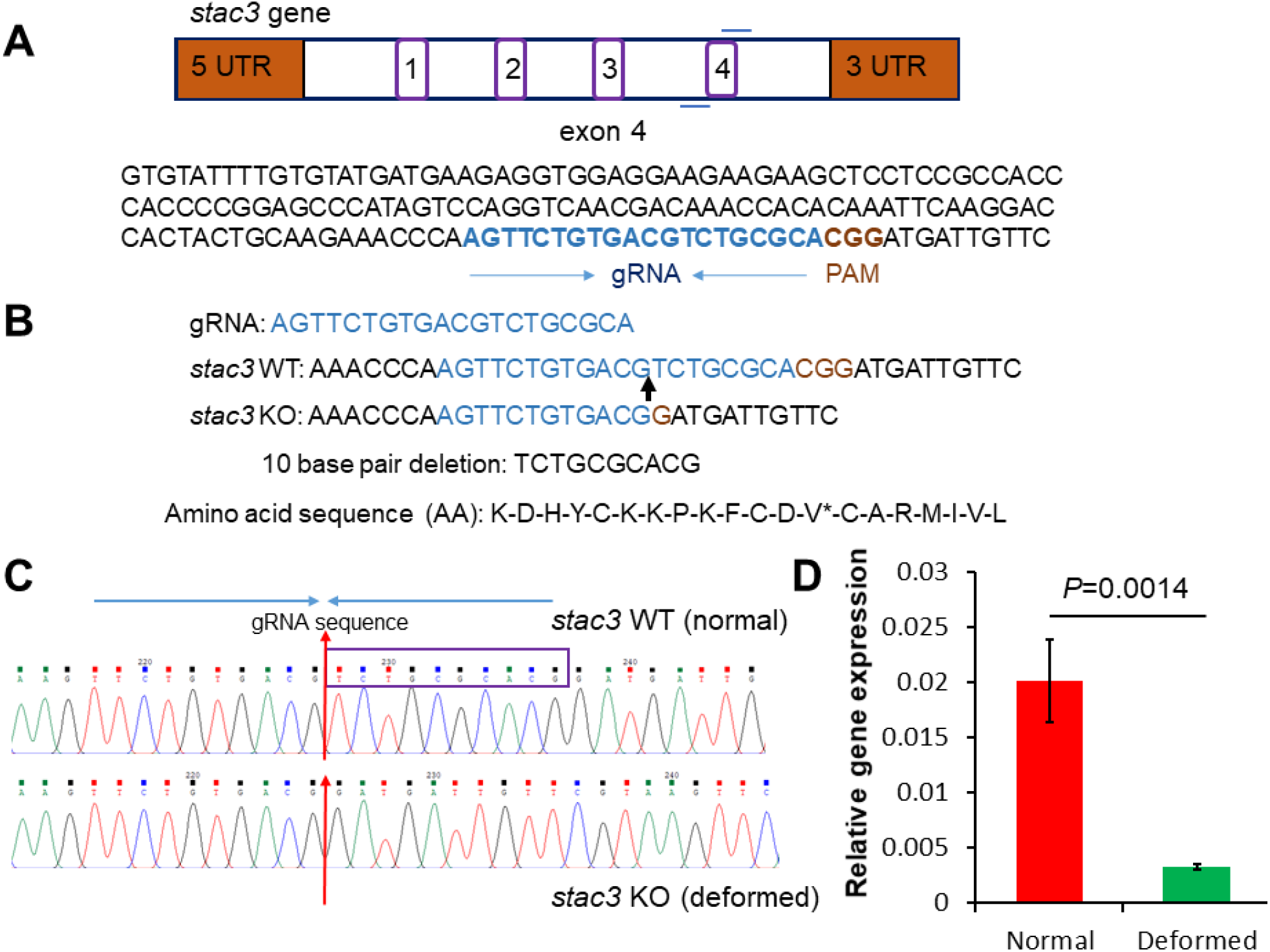
Deletion of 10 base pairs (bp) of *stac3* gene in zebrafish. (A, B). Schematic presentation of the *stac3* gene and nucleotide sequence of exon 4 region, which is marked for both guide RNA and PAM sequences, amino acids predicted using Ensembl.org. The stop codon at V78 is shown with asterisk. (C). DNA sequencing of the *stac3* wild type siblings (top), knockout (bottom). The deleted sequence TCTGCGCACG is marked by a violet box, red arrows indicate the starting position of the knockout. (D). RT-PCR analysis showed significant downregulation of *stac3* expression in deformed larvae (knockout) compared to wild type and heterozygous siblings (5 dpf). Data are presented as mean ± standard deviation. T-Test, *p= 0.0014.

### 2. *stac3* knockout constrains embryo hatching by altered Ca^2+^ level via *ryr1a* receptor in zebrafish

The *Stac3* gene plays a vital role in calcium homeostasis by modulating calcium-sensing dihydropyridine (DHPR) receptor and calcium-releasing ryanodine (RYR1) receptor at the sarcoplasmic reticulum^20^. We presumed that a mutation in *stac3* gene causes muscle weakness by altering physiological calcium level in zebrafish at early life. Intercrosses of adult *stac3*^+/-^ parents produced all three genotypes: *stac3^+/+^*, *stac3^+/-^* and *stac3^-/-^* larvae; the proportion of these genotypes complied with the Mendelian ratio. On 4 dpf, there was a significant (p<0.05) delay in hatching of stac3 knockout larvae (12%) compared to wild type and heterozygous control siblings (76.7%) (Fig2A, B). Under physiological conditions, muscle function (contraction) begins at 17 hours post-fertilization in ZF, such that, by 2 dpf, larvae are completely shed their chorion through rapid contractions^22^. Therefore, we sought to determine whether mutation in *stac3* alters lipid metabolism, resulting in damage to muscle fiber functional organization, consequently promoting the early death of ZF larvae.

**2.**
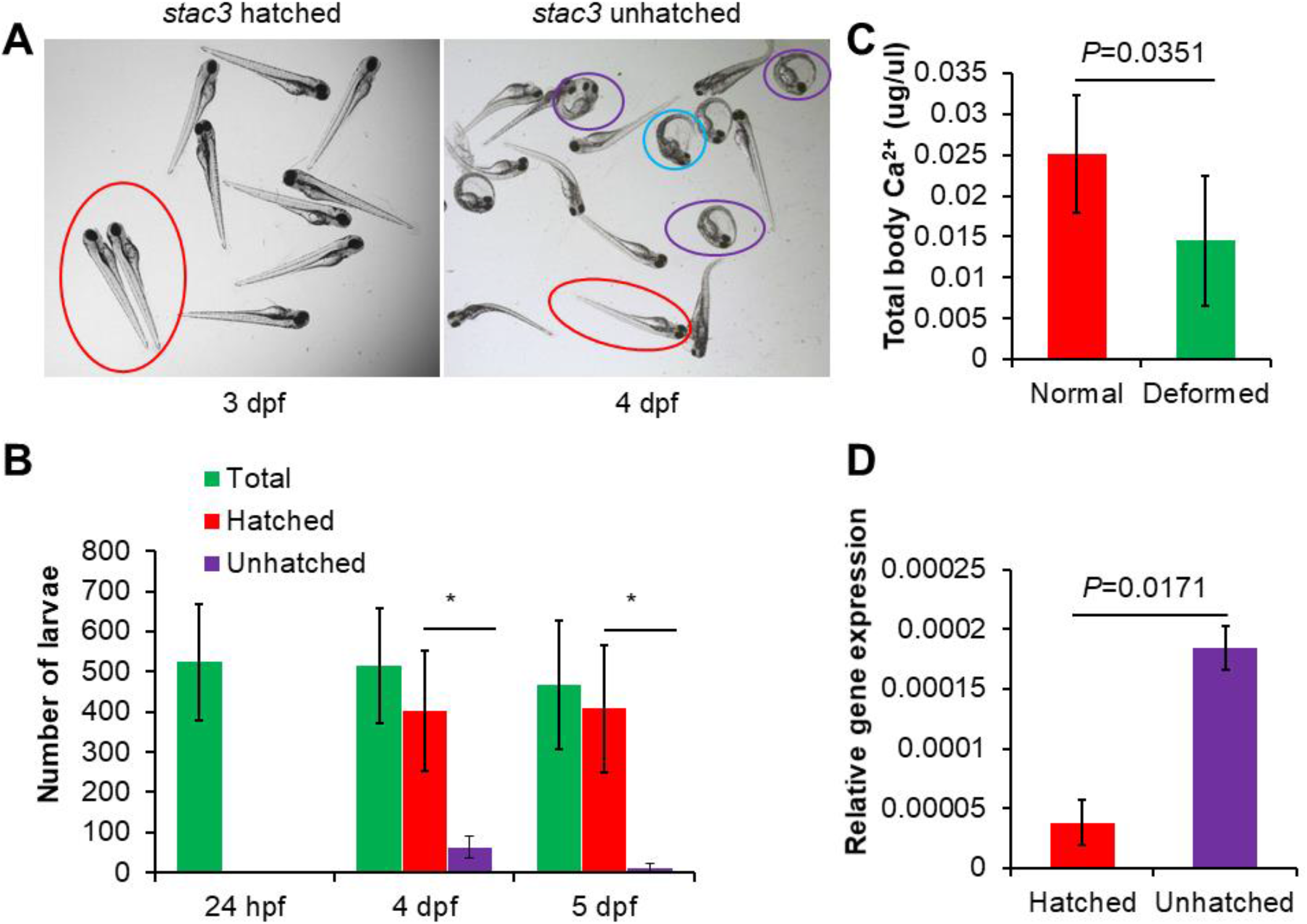
Delayed hatching of *stac3* ^-/-^ (KO) zebrafish larvae. (A). Hatching of *stac3^-/-^* larvae at 3 and 4 days post-fertilization (dpf). The blue circle marks larvae attempting to come out of the chorion (delayed hatching: knockout), while violet color indicates unhatched (knockout) larvae, hatched (*stac3*^+/+^ and *stac3*^+/-^: wild type and heterozygous siblings) larvae are outlined in red. (B). Percentage of delayed hatching of *stac3^-/-^* larvae at different time points, T-Test, *p=0.018 (4 dpf), *p=0.012 (5 dpf). (C). Whole-body Ca^2+^ levels in *stac3^-/-^* larvae, T-Test, *p=0.0351. (D). Ryanodine receptor 1a (*ryr1a*) gene expression, T-Test, *p=0.0171. Data are presented as mean ± standard deviation.

Indeed, we found that *stac3*^-/-^ larvae carrying 10 bp deletion mutation died within the chorion at 4 dpf, which was not seen in their controls siblings (FigS2A), suggesting that severity of declined muscle function (muscle weakness) gradually escalated. Moreover, genotypes of hatched and unhatched larvae were verified by DNA sequencing of a pooled sample (n=10) from each group (FigS2B, C). To determine whether low Ca^2+^ constrain hatching of ZF larvae, the progeny of *stac3^+/-^* parents was categorized into contrasting phenotypic groups for measuring whole body Ca^2+^ levels. Whole-body Ca^2+^ levels were compared between hatched vs. unhatched and normal vs. deformed larvae. As anticipated, total-body calcium levels were significantly lower (p< 0.05; Fig2C) in *stac3* knockout in comparison to *stac3^+/+^* and *stac3^+/-^* (control) siblings at the early muscle fibers growth and development. In parallel, we found that the relative gene expression of ryanodine receptor 1a (*ryr1a*) was significantly (p=0.0171; Fig2D) elevated in *stac3^-/-^* larvae compared to their controls. These results suggest that a mutation in *stac3* leads to a calcium deficiency via *ryr1a* receptor, which impedes skeletal muscle function, as demonstrated by delayed embryo hatching and declined locomotion (Fig7A, B and Fig9).

### 3. F-actin inadequacy and slow muscle fiber organization in *stac3^-/-^* larvae

stac3 knockdown has been shown to affect myofibrillar protein assembly^16^; we sought to determine whether *stac3* knockout can affect the cytoskeleton of skeletal muscle i.e., F-actin and slow myosin wirings. In humans, BA-D5 antibody was known to recognize one of the myosin MHC (cardiac β) isoforms^30^. Therefore, we performed slow muscle (anti-Myhc; F59) staining for visualization of all myosin isoforms in knockout *stac3*, heterozygous and wild type siblings at the age of 5 and 7 dpf. Consequently, disorganized slow muscle fibers and a deficit in conventional organization of individual fibers were identified in trunk and tail regions of *stac3^-/-^* larvae compared to *stac3^+/+^* and *stac3^+/-^* siblings at 5 dpf, this was more prominent with larger gaps at somite boundaries at 7 dpf (Fig3A-C and FigS3A and FigS4A, C panel). Furthermore, whole-body F-actin fibers were reduced in the *stac3* KO in contrast to *stac3^+/+^* and *stac3^+/-^* siblings (Fig3D and FigS3B). The myosin structural organization is essential for energy production of skeletal muscle for contractions^31^. In comparison to *stac3^+/+^* and *stac3^+/-^* control siblings, *stac3^-/-^* larvae showed a significant reduction in relative expression of both *srebf1* (p=0.0025) and its down-stream target acetyl co-enzyme-A (p=0.004), that are involved in lipogenesis (FigS3D). Therefore altered cytoskeleton architecture of skeletal muscle could impair metabolic processing of nutrients (lipids) in *stac3* KO.

**3.**
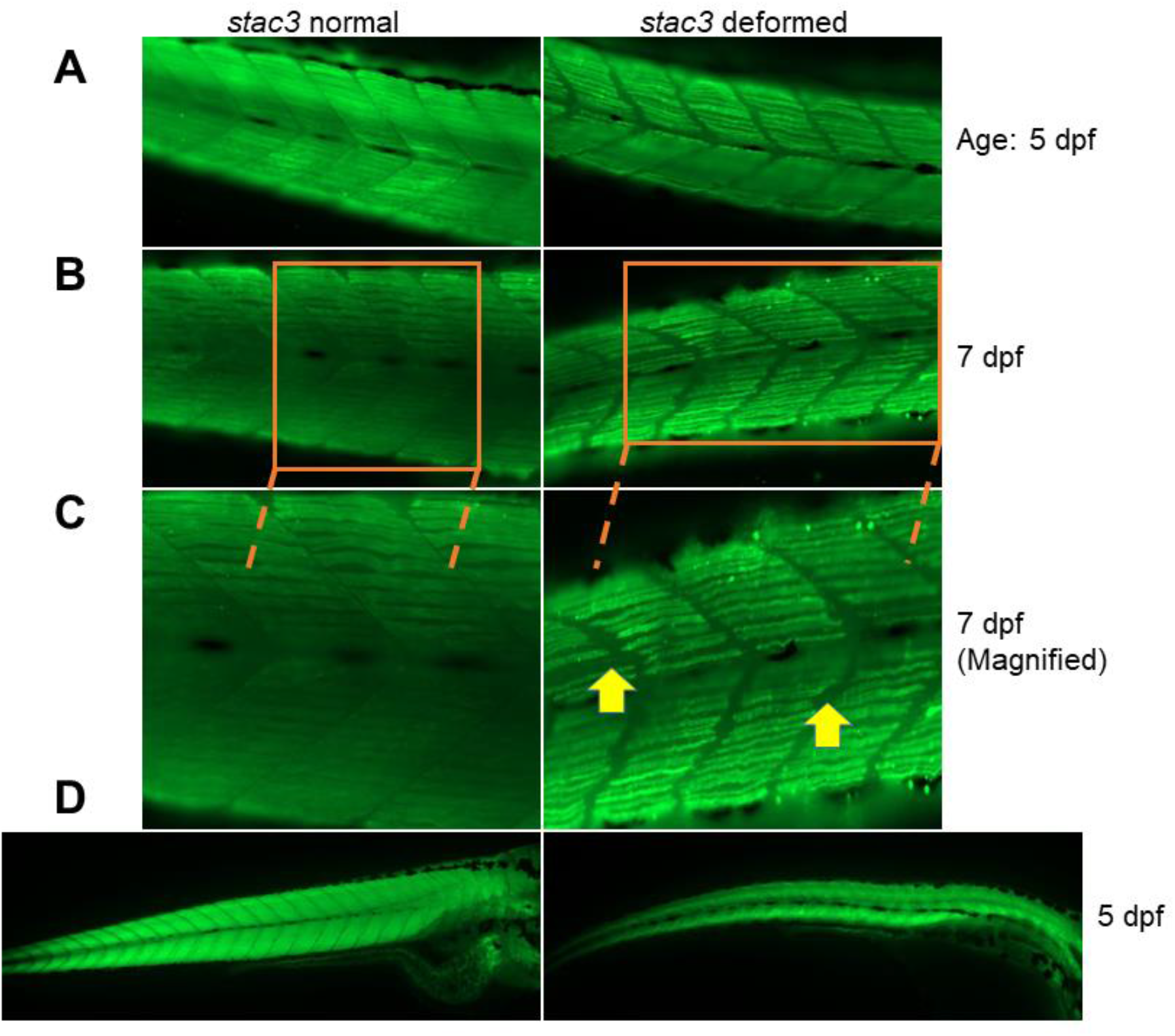
Slow muscle myosin staining of KO larvae. (A). Whole-mount immunostaining of slow muscle myosin isoforms with anti-Myhc1 antibody, in the trunk region, *stac3* KO larvae showed thin, curly muscle fibers, and smaller gaps compared to *stac3* WT and heterozygous fish at 5 dpf. (B). In the trunk region, *stac3* KO larvae (right) exhibited protruding breaks between somite boundaries compared to muscle fibers patterning’s in *stac3*^+/+^ and *stac3*^+/-^ control siblings (left) at 7 dpf. (C). Bottom panels (B) are magnified; protruding breaks between somite boundaries are marked with yellow arrow heads. (D). Wild type and heterozygous *stac3* and *stac3* knockout larvae stained with phalloidin at 5 dpf. Reduced amounts of filamentous (F-actin) fibers were noted in whole body of *stac3* knockout larvae (right) compared to *stac3*^+/+^ and *stac3*^+/-^ control siblings (left).

### 4. Expression of myogenic regulators is dysregulated in *stac3^-/-^* larva

*STAC3* is highly expressed in skeletal muscle tissue. Therefore, we sought to check whether the fragility of the musculoskeletal system in early life of the *stac3^-/-^* is dictated by impaired myogenesis, relative gene expression of myogenesis markers was evaluated. The lineage of skeletal muscle in the embryonic stage is determined by myogenic regulatory factors such as *MYOD*, *MYF5*, and *MYOG*^32^. Similarly, RT-qPCR analyses identified significant upregulation of *myoD* (p=0.001) alongside downregulation of *myf5* (p=0.03) and unaltered expression of *myoG* in *stac3^-/-^* larvae as compared to wild type and heterozygous controls at 5 dpf (Fig4A-C). Genetically, *MYOD* and *MYF5* are *bona fide* for myoblast fusion by maintaining the functional organization of actin and myosin fibers, while *MYOG* involves in the terminal differentiation of muscle cells^33^. Disorganized slow muscle fibers with a reduced amount of F-actin were found at 5 and 7 dpf in *stac3^-/-^* larvae (Fig3). However, gene expression of skeletal muscle markers such as *sox6*, slow muscle myosin (*smyhc1*), and troponin (*tnnca1*) were unchanged in *stac3^-/-^* compared to the *stac3^+/+^* and *stac3^+/-^* groups at 5 dpf (FigS3C). To our notice, it was shown that *MyoD^-/-^* mice are viable and fertile while contrastingly double knockout of *MyoD* and *Myf5* had led to a perinatal death^34^. Further, we show that unaltered *myoG* expression of *stac3^-/-^* larvae indicates defective signals that could commence in myogenesis before the formation of complete skeletal muscle fiber (Fig4D). Furthermore, abnormal histopathological features and compromised myogenesis were reported in myopathies^35^. Altogether, our results suggest that both *myf5* and *myoD* might have compensatory mechanisms of action in *stac3^-/-^* larvae, to secure cytoskeleton integrity by actin and myosin fibers formation.

**4.**
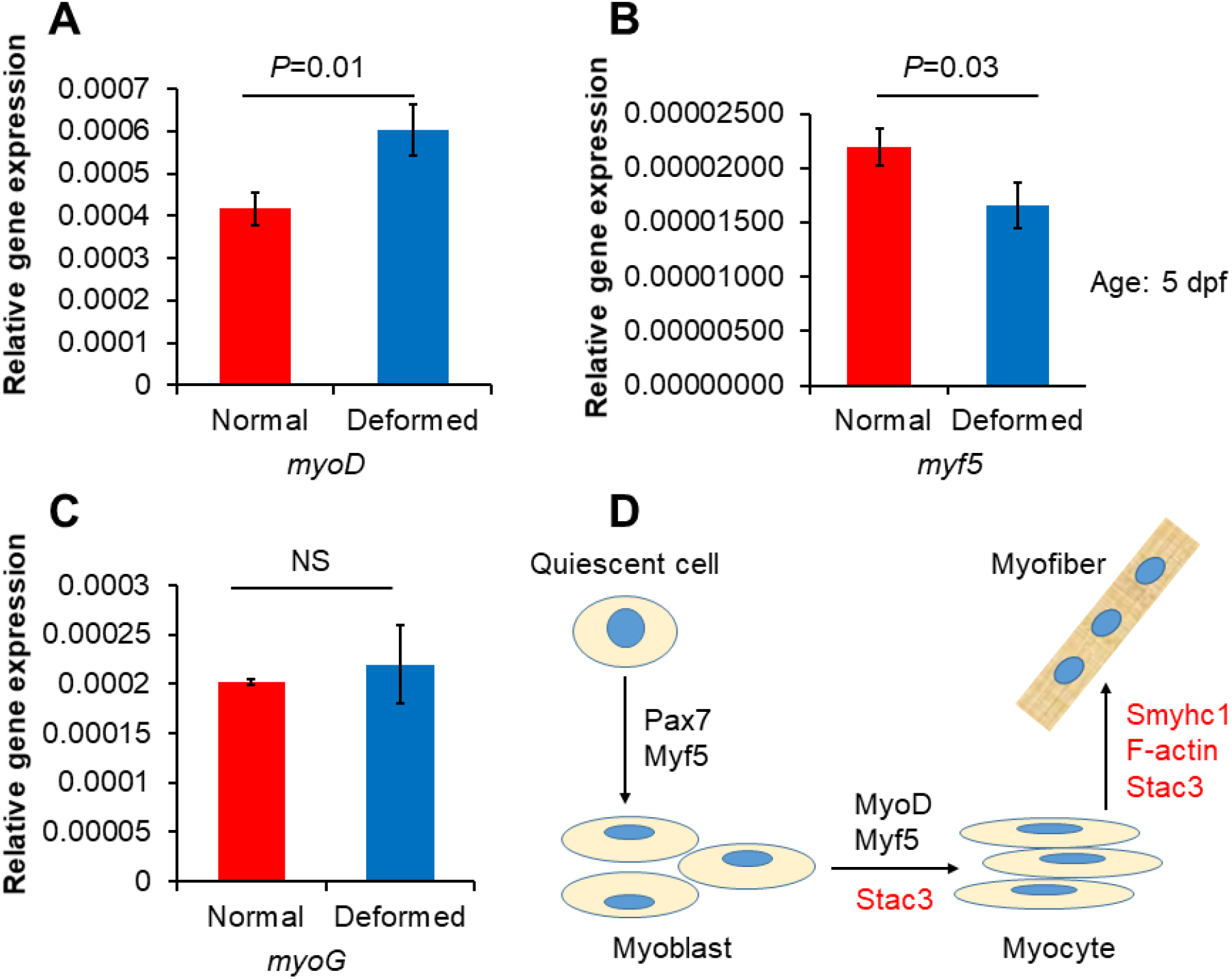
Knockout of *stac3* alters myogenesis in zebrafish larvae. (A). RT-PCR analysis revealed that gene expression of *myoD* was significantly upregulated in *stac3^-/-^* larvae compared to *stac3* wild type and heterozygotes (B). *myf5* gene expression was significantly downregulated in *stac3^-/-^* larvae compared to wild type and heterozygous siblings. (C). *stac3* knockout had no effect on mRNA levels of *myoG*. (D). A suggested mechanistic action of *stac3* in early muscle formation (red color denote functional regulators). T-Test, *p=0.01, *p=0.03, Non-significant (NS), n=10 animals for each group. Data are presented as mean ± standard deviation.

### 5. Distinct lipid metabolism in *stac3*^-/-^ fish at the early life

Besides in adipose and liver tissues, excess neutral lipids are often stored in skeletal muscle tissue, a hallmark of lipid storage myopathy. An increase in lipid droplets was revealed by electron microscope analysis of NAM patient skeletal tissue samples^12, 14^. Therefore, we sought to determine whether lipid metabolism is altered in *stac3*^-/-^ zebrafish and associated with impaired muscle fiber functional organization. At multiple developmental stages, *stac3* knockout and wild type and heterozygous siblings were stained with Oil Red O dye (ORO) for lipid visualization. In early life, zebrafish larvae utilize transported maternal lipids as the energy source for their active metabolic functions^36, 37^. Correspondingly, no differences in whole-body lipids of hatched (*stac3^+/+^* and *stac3^+/-^*) compared to unhatched (*stac3^-/-^*) larvae were observed on 4 dpf (Fig5A-A’). Notably, elevated neutral lipids were primarily observed in the yolk sac region of knockout larvae compared to *stac3^+/+^* and *stac3^+/-^* control siblings (p=0.0023; Fig5B-B’ and E) at 5 dpf, which corroborates the increase in lipid droplets measured in NAM patients^12, 38^. Impairments in genetic and metabolic process can affect lipid synthesis, transportation, utilization, and degradation in multiple cell and tissue types, and including skeletal muscle. We therefore further visualized neutral lipids of larvae by whole-mount ORO staining on the day of exogenous feeding (7 dpf) and, in knockout larva, found levels identical to those shown at 5 dpf (Fig5B-B’ and Fig5C-C’, D), suggesting that *stac3^-/-^* maintained surplus lipids, while in control groups these levels go down (FigS4B-B’). Of note, studies^16^ reported that *stac3* is involved in nutrition sensing, while its C1 domain binds to lipids^14^. Hence, we speculate that in presence of the *stac3* genetic background wild type and heterozygous siblings are capable of sensing nutrients for generating energy for performing enhanced muscle contractions, and survival of the larva (Fig7A, B and Fig9).

**5.**
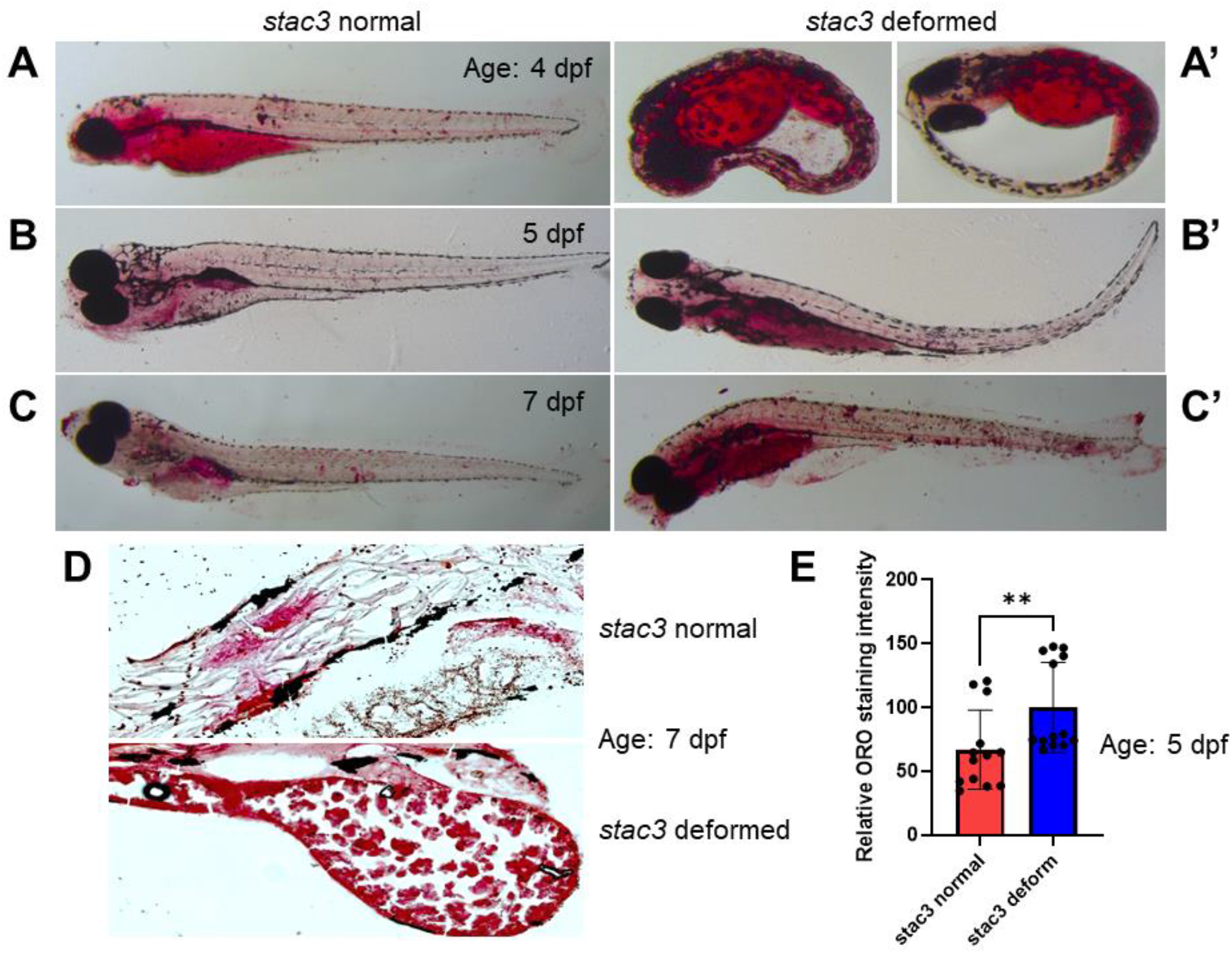
Lipid staining of *stac3* KO larvae using Oil Red O(ORO) (A-A’). Hatched (*stac3* wild type and heterozygous) and unhatched (*stac3* knockout) larvae were stained with ORO dye for neutral lipid visualization, both groups displayed similar levels of neutral lipids at 4 dpf (scale bar 3 mm). At 5 dpf (B-B’) and 7 dpf (C-C’: scale bar 3 mm), deformed (*stac3* knockout: B’ and C’) larvae exhibited higher neutral lipids compared to *stac3* wild type and heterozygous (B and C). (D). OCT based yolk sac sections of *stac3* normal and knockout larvae were stained with ORO dye. *stac3* knockout larvae (bottom) displayed more lipids compared to *stac3* normal (top) at 7 dpf. (E). Quantification red color indicated relative ORO staining intensity of the larvae. At each stage n=15 animals utilized per group. Mann Whitney test, **p=0.0023. Genotype of larvae (n=10) per group was confirmed as described in FigureS1A, B.

### 6. Metabolic dysfunction promotes death of *stac3* KO fish

The impact of accumulated lipids in NAM and pathophysiology of skeletal muscle tissue affected by this disease are unknown. However, detrimental effects on cytoskeletal elements and on their interactions in muscle tissue can lead to myopathies in humans^39^. We next asked whether a delay in embryo hatching, reduction in F-actin and alteration in slow muscle fibers organization weaken musculoskeletal system in *stac3* knockout. To confirm whether 10bp deletion mutation alters the structural integrity of *stac3^-/-^* skeletal muscle, birefringence analysis of *stac3^-/-^* found a patchy pattern of muscle fibers with dense black spots, whereas *stac3*^+/+^ and *stac3*^+/-^ control siblings had rich on muscle fibers with bright chevron structure on 6 dpf (Fig6A and TableS3). In this study, knockout larvae died gradually by 5 dpf, with obvious skeletal muscle defects; out of the expected by Mendelian ratio for *stac3* knockouts (25%) we found only ∼18.24% of knockout larvae survived until 7 dpf (TableS2). To understand the significance of genetic regulation of *stac3* in nutrients processing and its contribution to survival of zebrafish, we further maintained these surviving knockout larvae in an incubator and changed embryo water after feeding with nursery food, suggesting that *stac3* KO carries a systemic metabolic dysfunction manifested by increased neutral lipids (Fig5B, C panels). To determine whether high neutral lipid (nutrients load) levels can induce lethal lipotoxicity in *stac3* knockouts, we performed cellular senescence assay at the early development of zebrafish larvae^28^. As expected, our senescence-associated beta-gal probe (SA-β-gal) was strongly noticeable at the yolk sac region of the *stac3^-/-^* compared to *stac3^+/+^* and *stac3^+/-^* control siblings at 6 dpf (Fig6B, C and FigS5B, C panels). Furthermore, fasting of *stac3* KO larvae revealed a trend of improved survival between 7 and 10 dpf (p<0.0001; FigS6). These findings align with the lipid (ORO) staining observations and suggest that deletion of 10 bp in the *stac3* gene accelerates an early-age-specific senescence in cells, culminating in metabolic failure of the zebrafish larva (p<0.0001; Fig6D).

**6.**
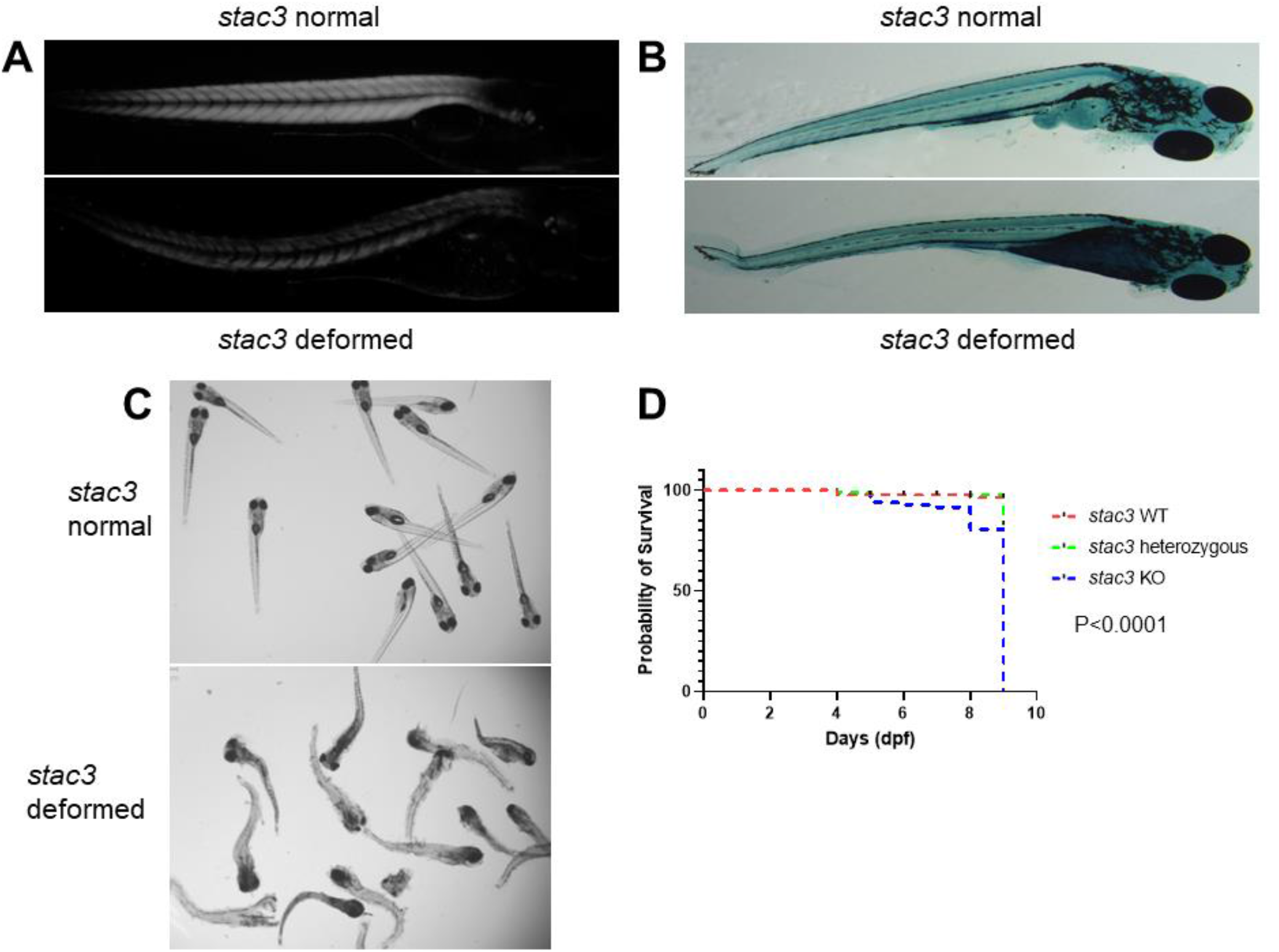
Death of *stac3* knockout might be due to metabolic dysfunction. (A). Birefringence assay demonstrated reduced chevron formation of muscle fibers in *stac3* knockout (bottom) compared to wild type *stac3* larvae (top) at 5 dpf. (B). *stac3* knockout larvae exhibited increased cellular senescence mostly in the yolk sac region (bottom) in comparison to wild type *stac3* (top), as demonstrated by SA-β-gal staining at 6 dpf. (C). Dead larvae are identified as *stac3* knockout (bottom) compared to *stac3* wild type and heterozygous siblings (top) at 5 dpf. (D). Survival analysis of *stac3^-/-^*, wild type and heterozygous siblings at 1-10 dpf. Post birefringence analysis, larvae (n=5-10) per group were utilized for genotype confirmation. Log-rank test applied for *stac3* WT, *stac3* heterozygous, and *stac3* KO zebrafish larva groups, ****p<0.0001. The genotype of dead larvae (n=80) was confirmed as described in FigureS1A, B.

**7.**
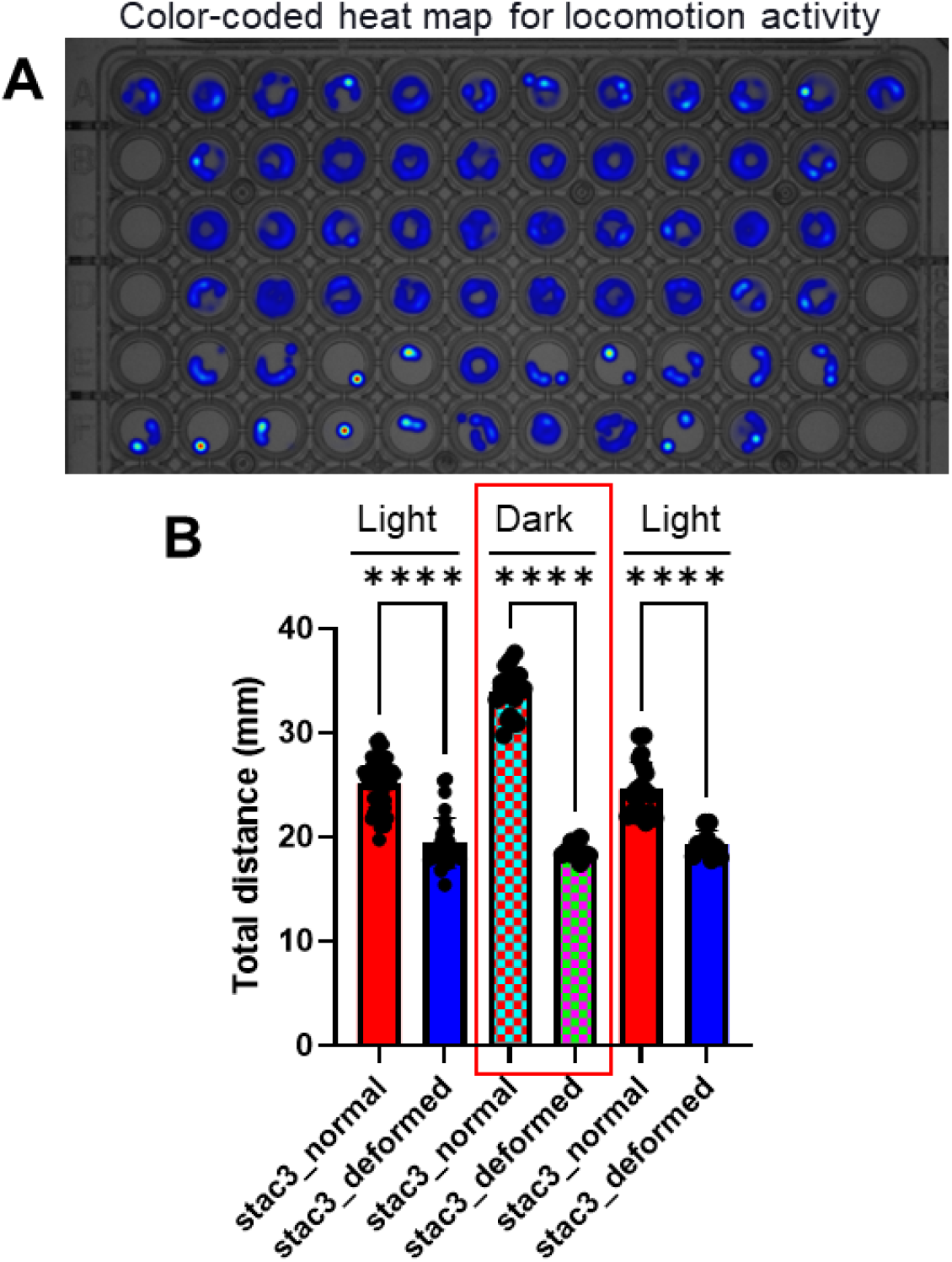
Swimming performance (locomotion) of *stac3* knockout in early life. (A). Merged, color-coded heat map for the total distance travelled by each larvae at 5 dpf. The blue color corresponds to a longer distance travelled, while pale yellow-red corresponds to no movement activity. (B). The locomotion tracking protocol included 15 minutes light on, 5 minutes light off, then 5 minutes light on. One-way ANOVA, ****p=0.0001, *stac3^+/+^* and *stac3^+/-^* (n=48) and *stac3^-/-^* (n=22) animals for each group. Genotype of larvae (n=10) per group was confirmed as described in FigureS1A, B. Each dot represents the total distance travelled by the larvae on graph Data are presented as mean ± standard deviation.

### 7. Caffeine treatment improves survival of the *stac3*^-/-^ larvae at 5 but not 7 dpf

We sought to determine whether caffeine could be an effective therapeutic molecule to a betterment of congenital myopathies due to its key mechanistic biochemical actions across muscle and other tissues^40^. Previous studies revealed that caffeine treatment had significantly enhanced muscle power with the improvement of Ca^2+^ release, increased fatty acid utilization, and decrease in the mortality of mice by antioxidant functions in brain^20, 41, 42^. We monitored the growth and survival of *stac3*^+/+^ and *stac3*^+/-^ (non-deformed) control siblings and *stac3*^-/-^ larvae with and without 5mM caffeine treatment at two different time points. After 5mM caffeine treatment for 96 hours, we discovered an improvement in the survival of the *stac3*^-/-^ larvae compared to untreated *stac3*^-/-^ control group at 5 dpf (Fig8A). Moreover, the difference of overall survival of the *stac3*^+/+^ and *stac3*^+/-^ control siblings was significantly (p<0.0001) increased compared to untreated and treated deformed animals 5 dpf. In the *stac3*^-/-^ zebrafish larvae at 7 dpf, we found that over 96 hours the *stac3*^-/-^ larvae were dying much faster (p<0.0001) than their untreated *stac3*^-/-^ and non-deformed control siblings (Fig8B). In contrast to 5 dpf, at 7 dpf deformed larvae did not show any improvement in their survival percentage after 5mM caffeine treatment compared untreated *stac3*^-/-^ (deformed) and treated *stac3*^+/+^ and *stac3*^+/-^ animals. In summary, the mechanistic regulation of *stac3* by caffeine is important during muscle fibers’ development at the early life (5 dpf), while caffeine regulation could be inhibited at the start of external feeding - in 7 dpf *stac3*^-/-^ zebrafish.

**8.**
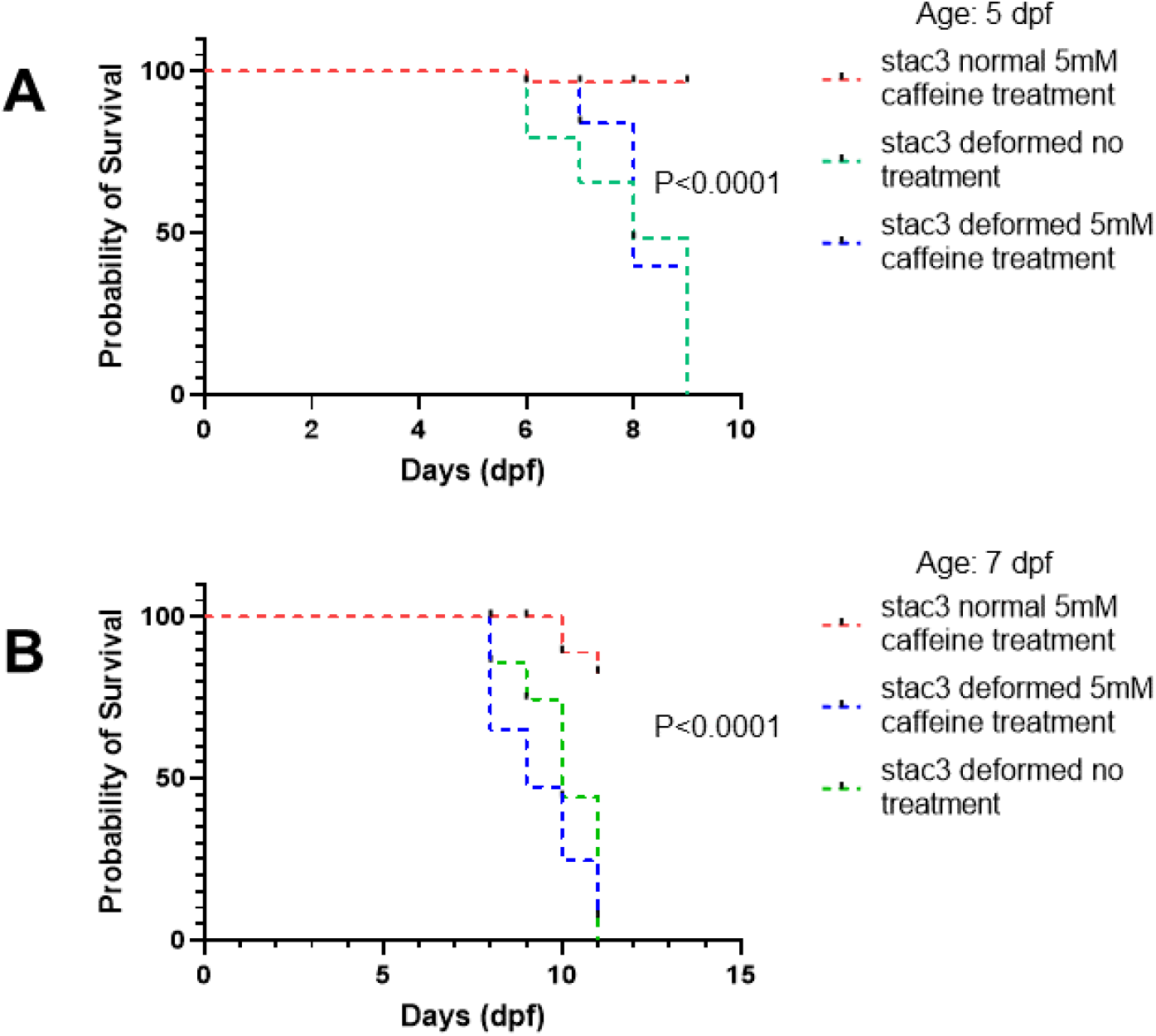
Survival of the *stac3*^-/-^ larvae at 5 dpf and 7 dpf after caffeine treatment. (A-B). Calculation of survival percentage of *stac3* KO (deformed) and *stac3* normal (*stac3^+/+^* and *stac3^+/-^* control siblings) zebrafish larvae with and without 5mM caffeine at 5 and 7 dpf. In both age groups, survival analysis was performed in 2-3 replicates, and larvae growth and survival were monitored for 96 hours. Log-rank test applied between treated and untreated deformed *stac3* zebrafish groups, ****p<0.0001.

**9.**
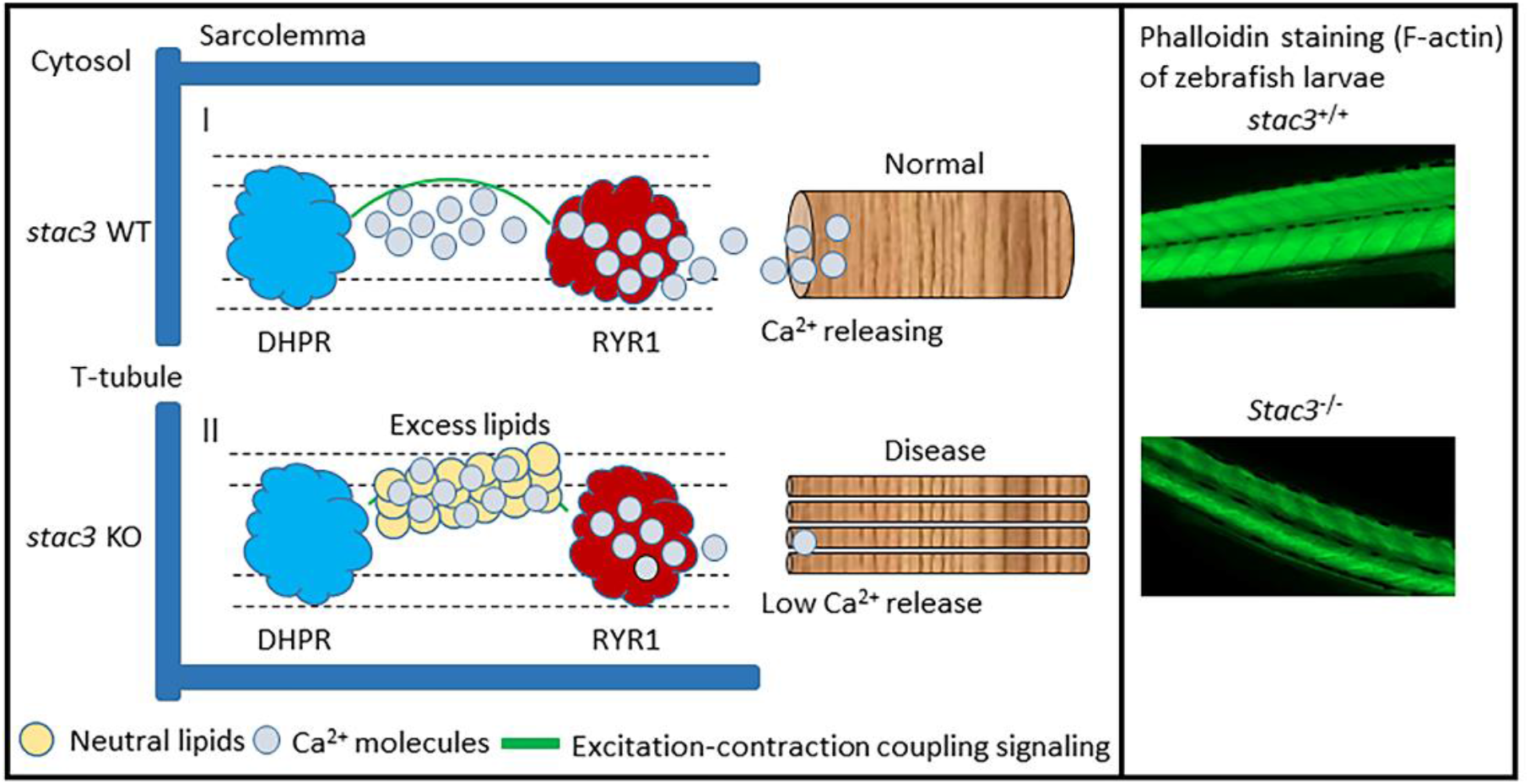
Working model of *stac3* gene in early life of zebrafish. In *stac3* knockout, increased lipids can directly reduce Ca^2+^ release at DHPR via RYR1 (*ryr1a*) receptor which seemingly delays the functional organization of F-actin and slow muscle fibers of zebrafish larva at early life.

## Discussion

In CRISPR/Cas9-generated *stac3* zebrafish mutants, we noticed that out of the expected Mendelian ratio of 25% only 18.24% *stac3* KO larvae survived till the age of 11 dpf. Additionally, functional and morphological observations of *stac3*^-/-^ larvae suggested muscle paralysis and skeletal muscle deterioration in the early life of *stac3^-/-^*. The present study thus explored the phenotypic basis for early life mortality of *stac3^-/-^* zebrafish and found that *stac3* is critical for early life lipid balance in ZF. Knocking-out this gene in zebrafish led to a delay in embryo hatching and to skeletal muscle defects, and eventually, to a gradual decline in survival at age 4-11 dpf. Under physiological conditions, muscle function (contraction) begins at 17 hours post-fertilization in ZF, such that, by 2 dpf, larvae should completely shed their chorion through rapid contractions. Of note, a proportion of *stac3*^-/-^ larvae died within the chorion (∼4 dpf), which is implying the existence of a paralyzed muscle function during early development. In CRISPR/Cas9 system-induced global knockout, we saw gross phenotypes early on, thus we identified larvae as normal (*stac3* wild type and heterozygous siblings) and deformed (*stac3* knockout) in the entire study. Earlier studies found altered ratios of slow and fast muscle fibers in *Stac3* KO mice, possibly reflecting impaired muscle metabolism^15, 43^. Zebrafish carrying a missense mutation of the *stac3* gene were shown to have EC coupling alteration-instigated swimming impairments, which preceded larval death^22^.

To improve the survival of the larvae beyond the age of 11 dpf, we applied special conditions (i.e.nursery feeding and replenishing with a fresh system water) to these 18.24% *stac3* KO larvae that survived, along with *stac3^+/+^* and *stac3^+/-^* control siblings. We thus discovered that between 7-11 dpf, expedited mortality was observed in *stac3^-/-^* fish only. Furthermore, fasting revealed a trend in better survival of fasted vs. fed *stac3* KO larvae between 7 and 10 dpf (FigS6). It could be that nutrient-dependent transcriptional regulation in *stac3* KO zebrafish failed in coping with exogenous feeding. We further sought to measure the expression of myogenic regulators and lipogenesis synthesis genes to correlate with metabolic activities involved in developing muscle fiber architecture. We found that loss of *stac3* significantly upregulated *myoD* and downregulated *myf5* expression, while *myoG* gene expression was unaltered, whereas *srebf1* and acetyl co-enzyme were significantly down-regulated, which suggests that myogenesis can be independently controlled by *myoD* and *myf5* genes^44, 45^. Congruently, *myoG* mutant larvae showed no effect on muscle phenotypes^45^. Of note, in chicken development, distinct diet conditions associated with increased mRNA levels of myogenic regulators (*Myf5*, *MyoD*, and *Myf4*) controlled the growth of skeletal muscle^46, 47, 48^. Recently, the expression of *MYF5* was reported to be significantly altered in human congenital myopathy^49^. In sum, transcription of the myogenic regulatory factors *myoD* and *myf5*, which encode proteins critical for fine-tuning of musculoskeletal health in zebrafish, is dependent on *stac3*.

In humans, clinical features of congenital myopathies include muscle weakness, increased intramuscular fat and muscle fiber degeneration^50^. Histologically and molecularly, congenital myopathies are categorized into sub-entities based on the affected protein (e.g. desmin, dystrophin, collagen) and lipid contents in skeletal muscle tissue^51, 52, 53^. Of note, NAM patients demonstrate muscle weakness and increased lipid levels^12, 54^. In our analysis of the gross appearance and organization of *stac3*^-/-^ larvae muscle fibers, *stac3* was found to play a pivotal role in establishing F-actin and slow muscle fiber functional coordination to generate strength and execute active contractions. Despite *Stac3* nutrient-sensing activity, its potential functional role via lipid metabolism has barely been studied in animal models for NAM. Our studies found that deletion of 10 base pairs in *stac3* had no impact on neutral lipid levels in 4 dpf *stac3^-/-^* fish. Yet, higher levels of neutral lipids were measured in *stac3^-/-^* compared to *stac3^+/+^* and *stac3^+/-^* control larvae on 5 dpf and maintained through 7 dpf. Taken together, these findings suggest that *stac3* acts as an age-specific metabolic regulator in the transformation from unhatched to hatched zebrafish larvae, by controlling lipid (energy) utilization in early life. Five dpf is an ideal age for investigating skeletal muscle formation while the effect of metabolic changes on muscle fibers organization can be examined at 7 pf in zebrafish. We observed that the damaged organization of F-actin and slow muscle fibers could play a role in the progression of myopathy. Due to a significant role in energy processing, utilization, and generating strength in humans, we suggest that improvement of slow muscle fiber integrity could delay the progression of muscle diseases (congenital myopathies)^55^.

According to the literature^56, 57^, the yolk is considered a reservoir of metabolically active nutrients needed for the embryonic growth and development of an organism. Around 3 dpf, yolk sac contains vascular network with compartments delimited by veins, while reduction of yolk sac is seen as embryo utilizes it at 4 dpf^37, 58^. Inherently, zebrafish larvae utilize maternally acquired lipids as an energy source for the routine biological functions until the age of ∼5 dpf, while the larva transforms into adult zebrafish with rapid development of musculoskeletal system^36, 59^. The yolk syncytial layer (YSL) is important for the delivery of lipids during the larval growth and development^58^. With functional activity in early-life stages and nutrients’ demand in an organism, lipids are essential for formation of organ and tissue types e.g. heart, liver, intestine, pancreas, and other vascularized regions, including skeletal muscle. Recently, it was reported that intramuscular adipose tissue is increased in dystrophy and neuromuscular disease patients compared with control individuals^60^; it now is evident that NAM’s etiology is similar. Altogether, the persistence of altered metabolic condition, which can deteriorate muscle function in NAM, is under-appreciated. We found that *stac3* KO larvae retained neutral lipids, mostly at the yolk sac region, even at the age when the exogenous feeding becomes necessary (7 dpf). At this age, neutral lipids were less visible in *stac3*^+/+^ and *stac3*^+/-^ control groups. Consequently, we speculate that the intact *stac3* enables nutrient sensing for growth of the musculoskeletal system, resulting in rapid muscle contractions, and larvae survival. Loss of *stac3* manifests by accumulation of neutral lipids, which is detrimental in the early growth stages of zebrafish larvae^61^. Storage of neutral lipids might contribute to a systemic metabolic dysfunction by lipotoxicity/senescence effect on cytoskeleton organization of the larvae. Of interest, the family of STAC proteins has been associated with nuclear factor-*k*B and C/EBP in controlling senescence in the course of muscle cell proliferation and differentiation^16^. We found that senescence-associated beta-gal staining was accentuated in the yolk sac region of *stac3^-/-^* at 6 dpf. Taken together, at early or late-embryonic stages, skeletal muscle cells might encounter lipotoxicity-induced stress, which triggers their dormant state through cellular senescence^31^. Prolonged accumulation of lipids via lipotoxicity in *stac3* knockout zebrafish could further contribute to unfavorable conditions (signals) that cause tissue (i.e. muscle) deterioration and larval death. Comprehensive studies will be required to understand lipid metabolism-driven vertebrate developmental transformations in early ZF life.

Physiologically, STAC3 oversees the biochemical relationship between calcium sensing (DHPR) and calcium-releasing (RYR1) receptors at the sarcoplasmic reticulum in skeletal muscle^20, 43^, with calcium homeostasis being essential for active contractions, tissue formation, maturation, and regeneration^62^. In the present work, *stac3*^-/-^ larvae were found to have lower concentrations of calcium and upregulation of *ryr1a* in an early patterning of skeletal muscle fibers. These findings compelled us to draw a working hypothesis that escalated lipid levels can directly reduce muscle contractions, as manifested by (a) a delay of embryo hatching due to muscle weakness, (b) impaired Ca^2+^ release via *ryr1a*, and (c) damaged organization of F-actin and slow muscle fibers (Fig9). In sum, these *stac3* KO phenotypes suggest paralyzed muscle function manifested as a significantly reduced locomotion on 5 dpf. Recent case studies from Turkey^63^, France^64^ and Russia^65^ suggested that pathophysiology of myopathy progression is comparable to NAM disease phenotypic outcome in humans. 36% of individuals afflicted with Native American myopathy (NAM) died by the age of 18 years^12^. We hypothesized that caffeine could be an effective therapeutic molecule to a betterment of congenital myopathies due to its key mechanistic biochemical actions across muscle and other tissues^20, 41, 42^. Indeed, our findings suggest that there is increased trend in the survival of the 5mM caffeine treatment in deformed *stac3* larvae at 5 dpf, while 5mM caffeine treatment revealed rather an opposite effect on *stac3*^-/-^ at 7 dpf. Hypothetically, the biochemical action of caffeine could be inhibited at the age of external feeding should start, which is ∼7 dpf. Thus, we postulate that our *stac3* knockout zebrafish, established for NAM is a lipid-related human congenital myopathy (lipid storage myopathy). Altogether, exploring early life’s genetic mechanisms using *stac3*^-/-^ zebrafish model are frontiers for repressing muscle fiber degeneration and refining muscle function maintaining lipid homeostasis in NAM disease during the course of muscle fibers formation.

Animal models are the gold standard for gaining pre-clinical insights into various diseases, due to their anatomical and morphological resemblance to humans^66^. Of late, modeling of human myopathies is achieved by knocking out disease-causing genes by CRISPR/Cas9 technology^67^. The current work demonstrated that *stac3^-/-^* fish may serve as a potential model for identifying lipid-based biomarkers or small molecules for the early diagnosis or treatment of NAM. Intramuscular adipose tissue is increased in dystrophy and neuromuscular disease patients compared with control individuals. We suggest that *STAC3* has beneficial effects on muscle metabolism by preventing the damage triggered by genetic or metabolic dysfunctions due to abnormally accumulated lipids.

## Acknowledgements

We are grateful to Dr. Chen Shochat for her helpful advice in CRISPR. The authors would like to thank Dr. Sergio Szvalb’s lab at the Department of the Pathology, Ziv Hospital, Safed, Israel, for the muscle histology. We extend our gratitude to Dr. Daya’s lab for allowing us to perform the birefringence assay at the Faculty of Marine Sciences, Ruppin Academic Center, Mikhmoret, Israel.

## Data availability

Data generated or utilized in this study can be found in online data repository source.

## Supplementary materials

The supplementary materials and figures related to this article can be accessed through online version of the manuscript.

## Author Contributions

RD: Conceptualization, investigation, data curation, data analysis, writing original draft, manuscript review and editing, DK: Conceptualization, data curation, funding acquisition, project administration, supervision, manuscript review and editing. HFZ: investigation, data curation, data analysis and manuscript review and editing. All the authors contributed to the article and approved the version submitted for the publication.

## Supplementary materials and methods

With the guide RNA targeted exon-4 sequence of *stac3* gene, we identify suitable restriction enzymes using NEB Cutter (https://nc3.neb.com/NEBcutter/) for genotyping zebrafish larvae, while the guide RNA sequence has a restriction digestion site at its 3 prime for genotyping.

### stac3

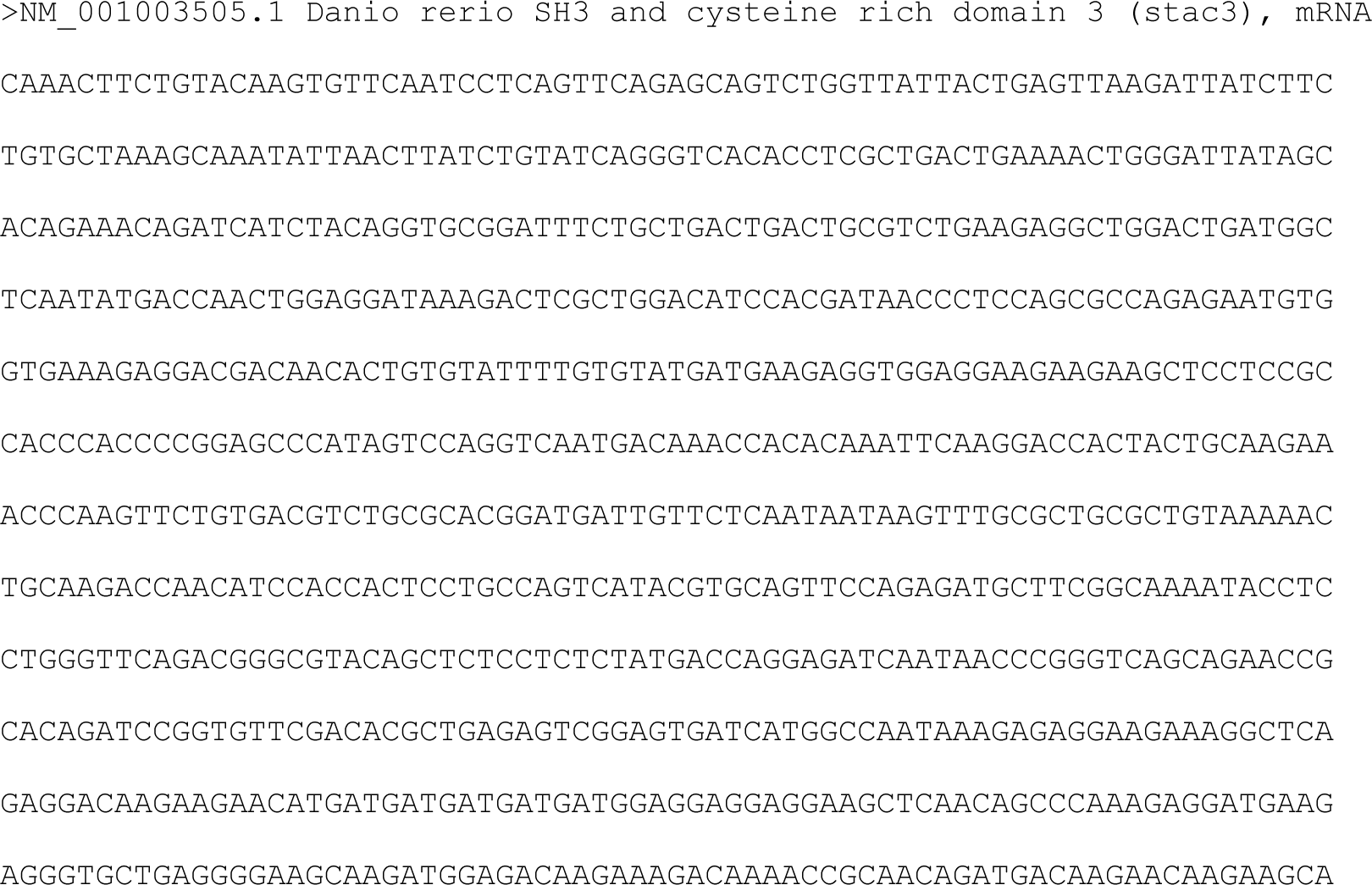
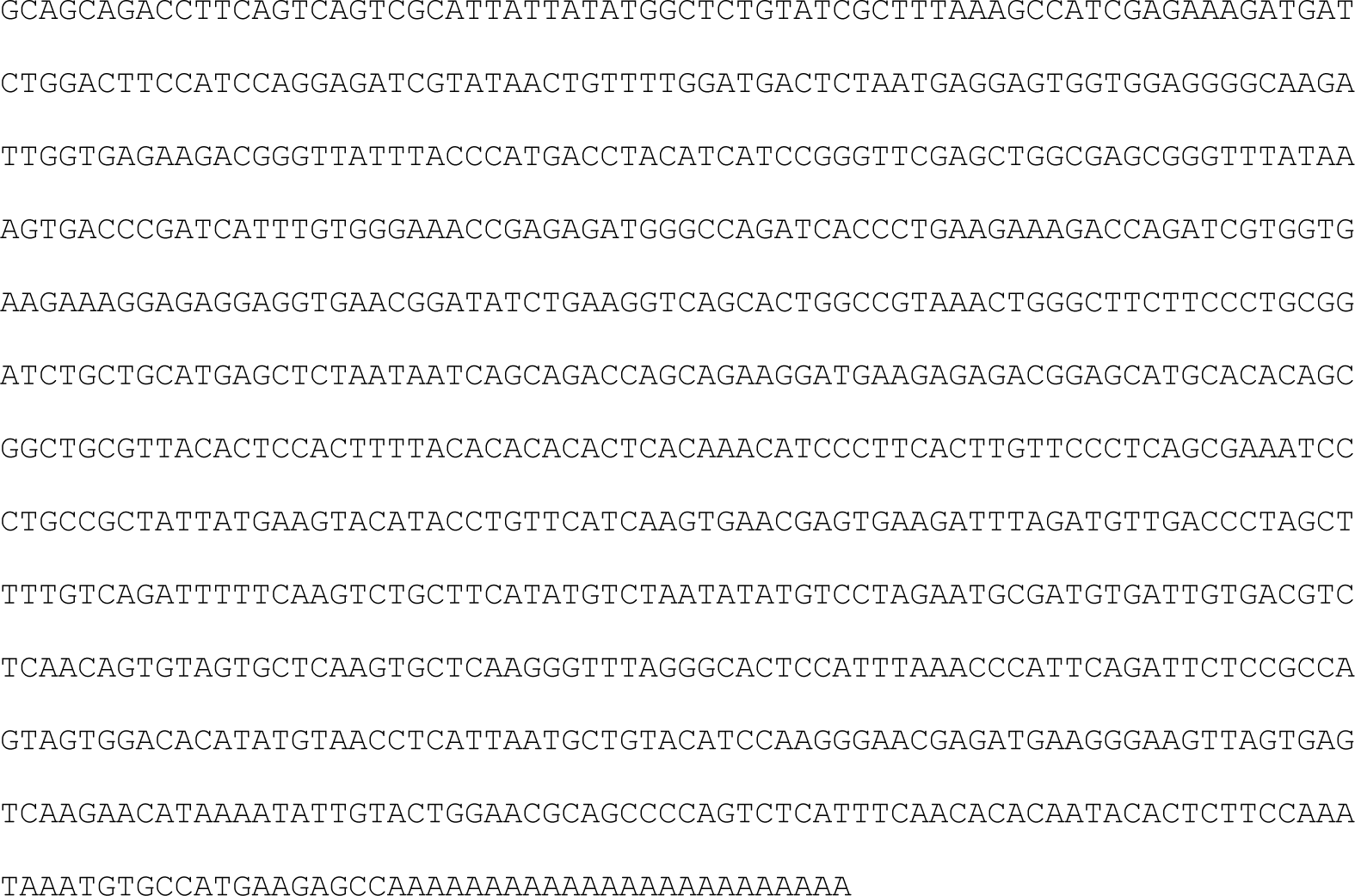

### gRNA

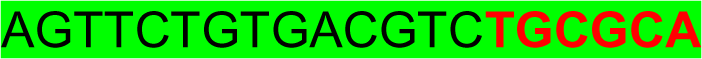

### *stac3* exon 4

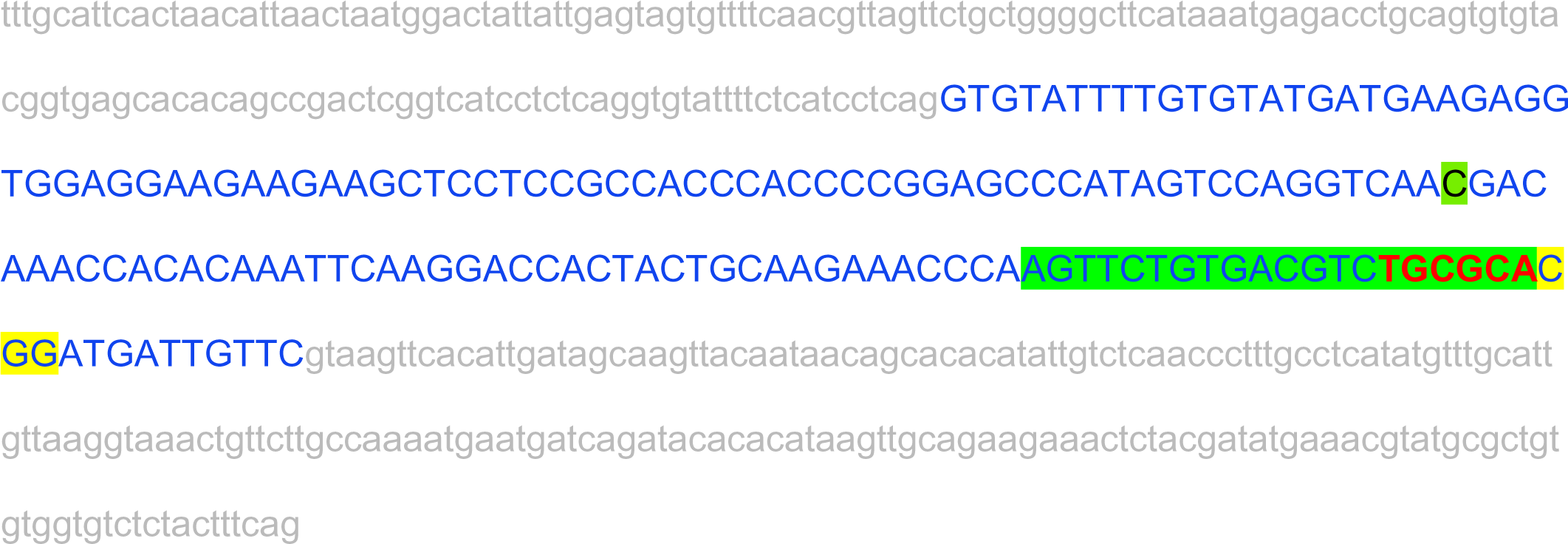

FspI restriction enzyme

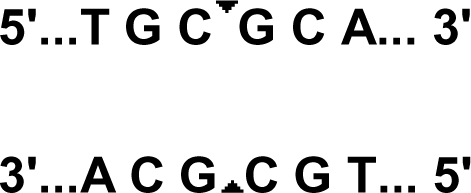

Pcr product 363bp

After cut 247bp 113bp

Genotyping of *stac3*^-/-^ vs. *stac3*^+/+^ and *stac3*^+/-^ control siblings was confirmed as described in FigureS1A, B. Blue: coding region, green color marks the gRNA region in exon4 of the *stac3* gene and protospacer adjacent motif sequence is in yellow, while red denotes restriction digestion site.

## Supplementary figures and legends

**S1.**
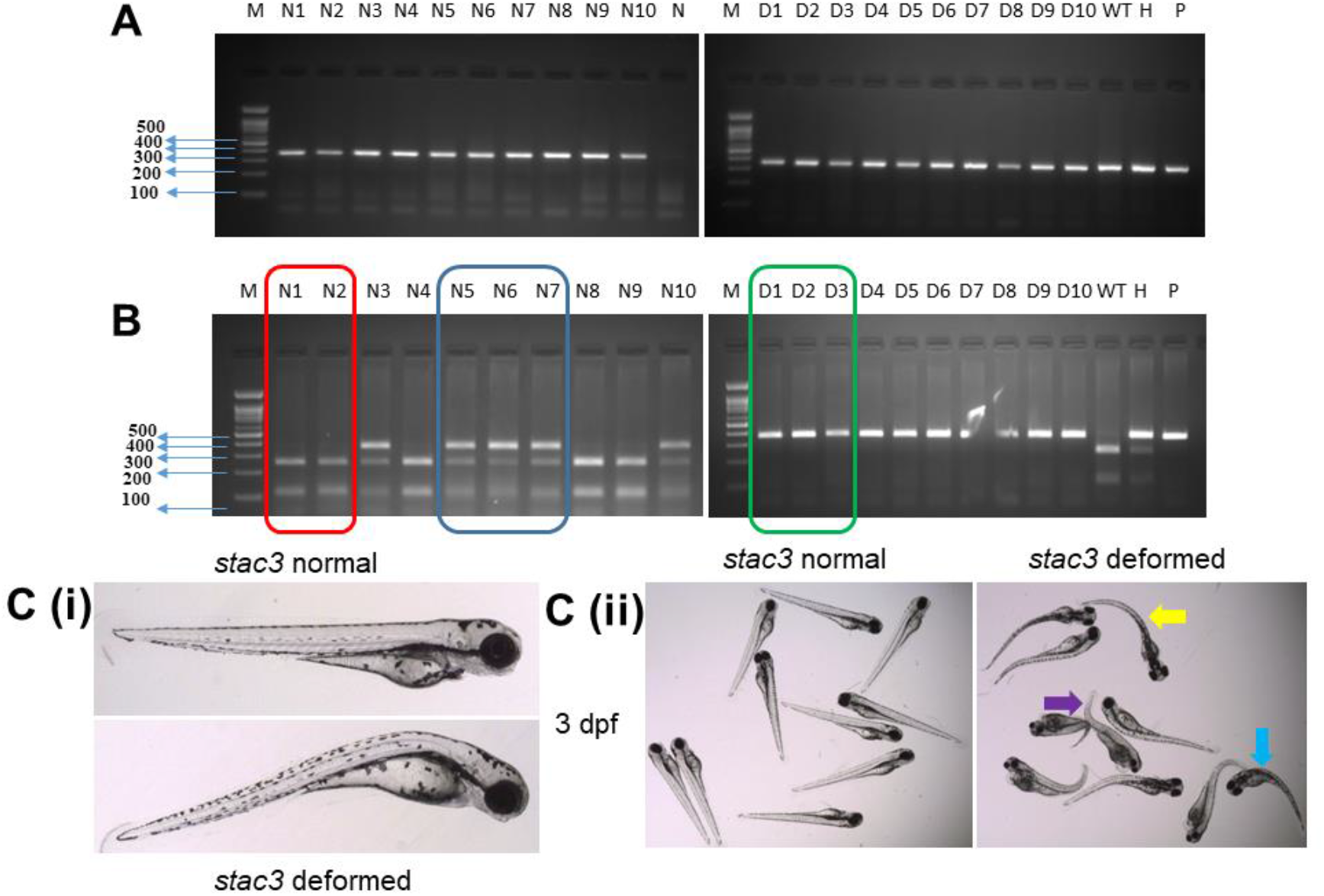
Knockout confirmation of the *stac3* gene in zebrafish. (A). DNA was extracted from wild type and heterozygous and knockout *stac3* larvae at the age of 3 dpf and amplified in a polymerase chain reaction (PCR) using *stac3* gene-specific primers (top). (B). The PCR product was confirmed on a 2% agarose gel, which showed a single band at 363 base pairs (bp). The PCR product was digested using the *FSPI* enzyme. The wild type had two digested DNA products (250 bp and 113 bp, red). The heterozygotes had three digested DNA products (113 bp, 250bp, and 363 bp, blue). The knockout undigested PCR product was a single band at 363 bp (green) (bottom). M-marker, N1-N10 (normal: *stac3^+/+^* and *stac3^+/-^* siblings): wild type and heterozygous, D1-D10 (deformed: *stac3^-/-^*): knockout, WT-wild type, H-heterozygote, P-positive control. (C). At 3 days post-fertilization, *stac3^-/-^* mutant larvae were distinguishable from *stac3^+/+^* and *stac3^+/-^* siblings (i), and displayed multiple congenital musculoskeletal defects, which included bending at the head (blue), trunk (yellow), and tail regions (violet) (ii). Scale bar: 1 mm.

**S2.**
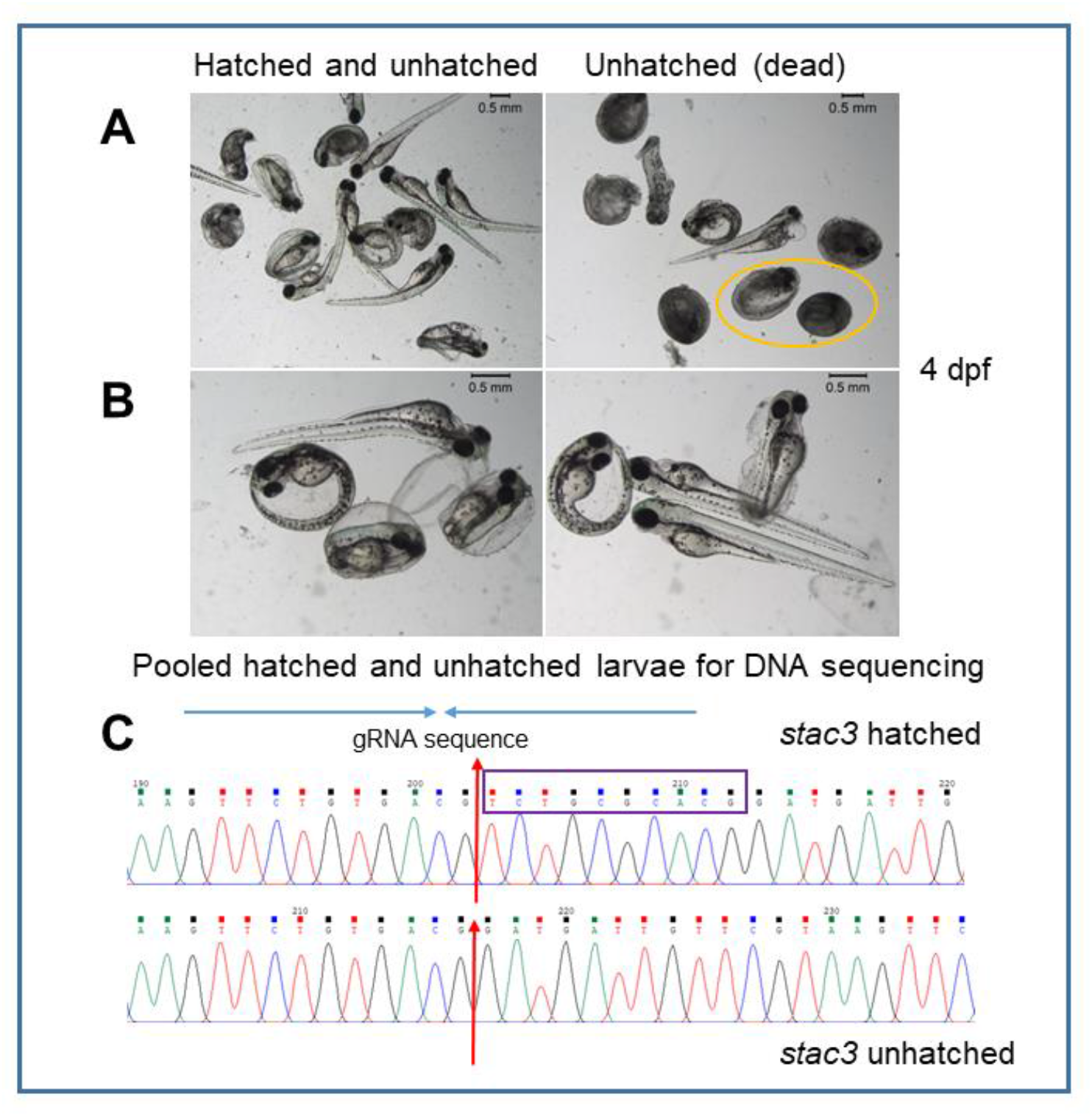
Genotype confirmation of delayed hatching of larvae by sequencing at 4 dpf. (A). Hatched and unhatched *stac3* larvae (left panel). Dead *stac3^-/-^* larvae within chorion are shown in the yellow circle in the right panel. (B). Representative images of hatched and unhatched *stac3* larvae (n=10) pooled for sequencing (left and right panel). (C). All hatched and unhatched larvae confirmed as wild type (top), heterozygous (not shown), and knockout larvae (bottom) respectively. Knockout larvae acquired a 10bp deletion mutation, the TCTGCGCACG sequence is marked by a violet box. Scale bar: 0.5 mm.

**S3.**
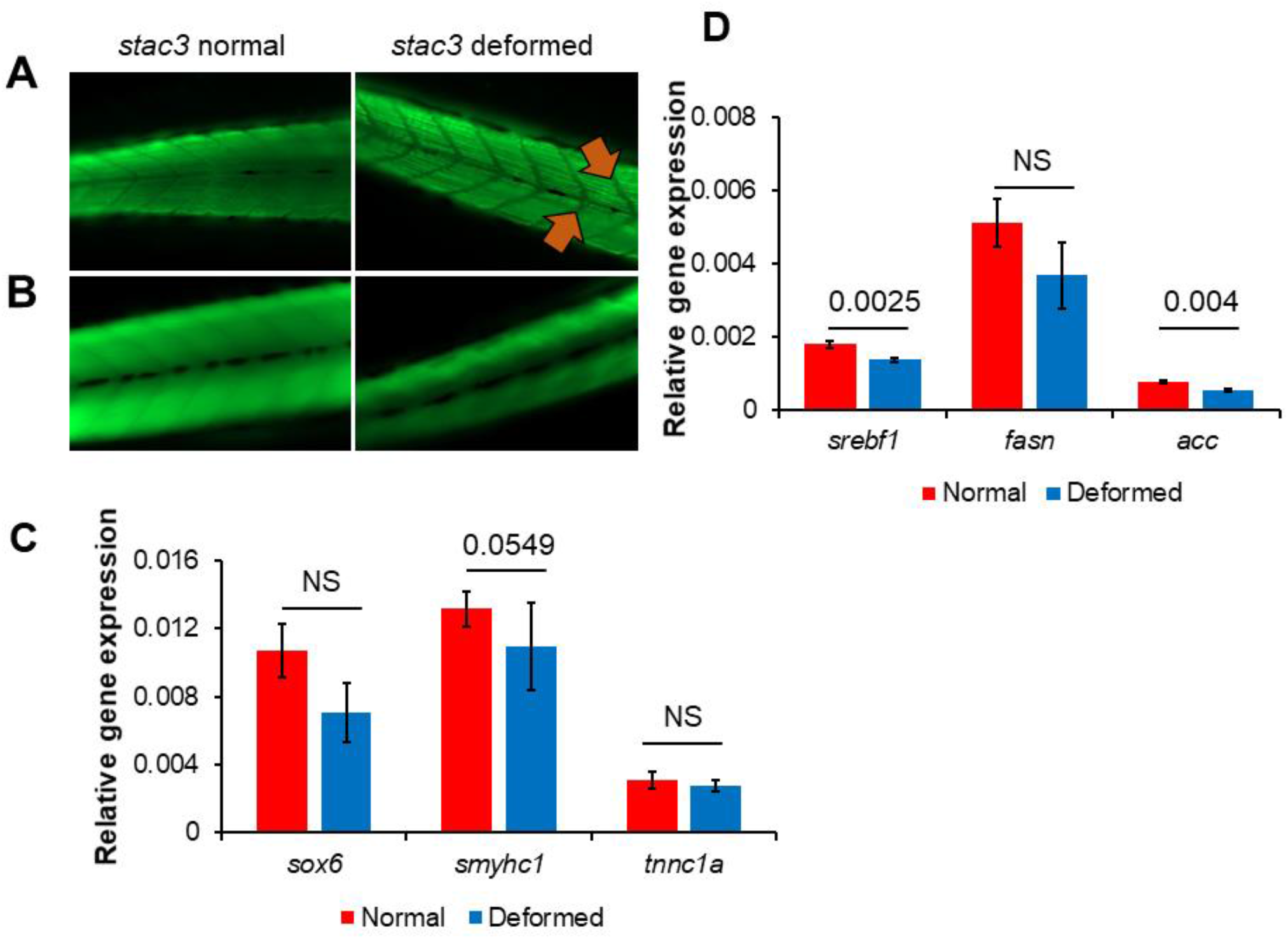
F-actin and slow muscle myosin staining of knockout larvae. (A). Tail region of wild type and heterozygous *stac3* and *stac3* knockout larvae stained with F59 antibody. *stac3^-/-^* larvae (right) showed disorganized slow muscle fibers compared to *stac3* wild type and heterozygous siblings (left), while orange arrowheads indicate alterations in muscle fibers. (B). Reduced amounts of filamentous (F-actin) fibers were noted in whole body of *stac3* knockout larvae (right) compared to control siblings (left) stained with phalloidin and followed by ‘20X’ magnification at 5 dpf. (C). RT-PCR analysis revealed that gene expression of skeletal muscle markers such as *sox6, smyhc1*, and *tnnca1* was unaltered, (D). whereas *srebf1* and its down-stream target acetyl co-enzyme-A were significantly down regulated, in *stac3^-/-^* larvae compared to *stac3* “normal” (wild type and heterozygotes). T-Test, **p=0.0025 and p=0.004. Non-significant (NS), n=10 animals for each group. Data are presented as mean ± standard deviation

**S4.**
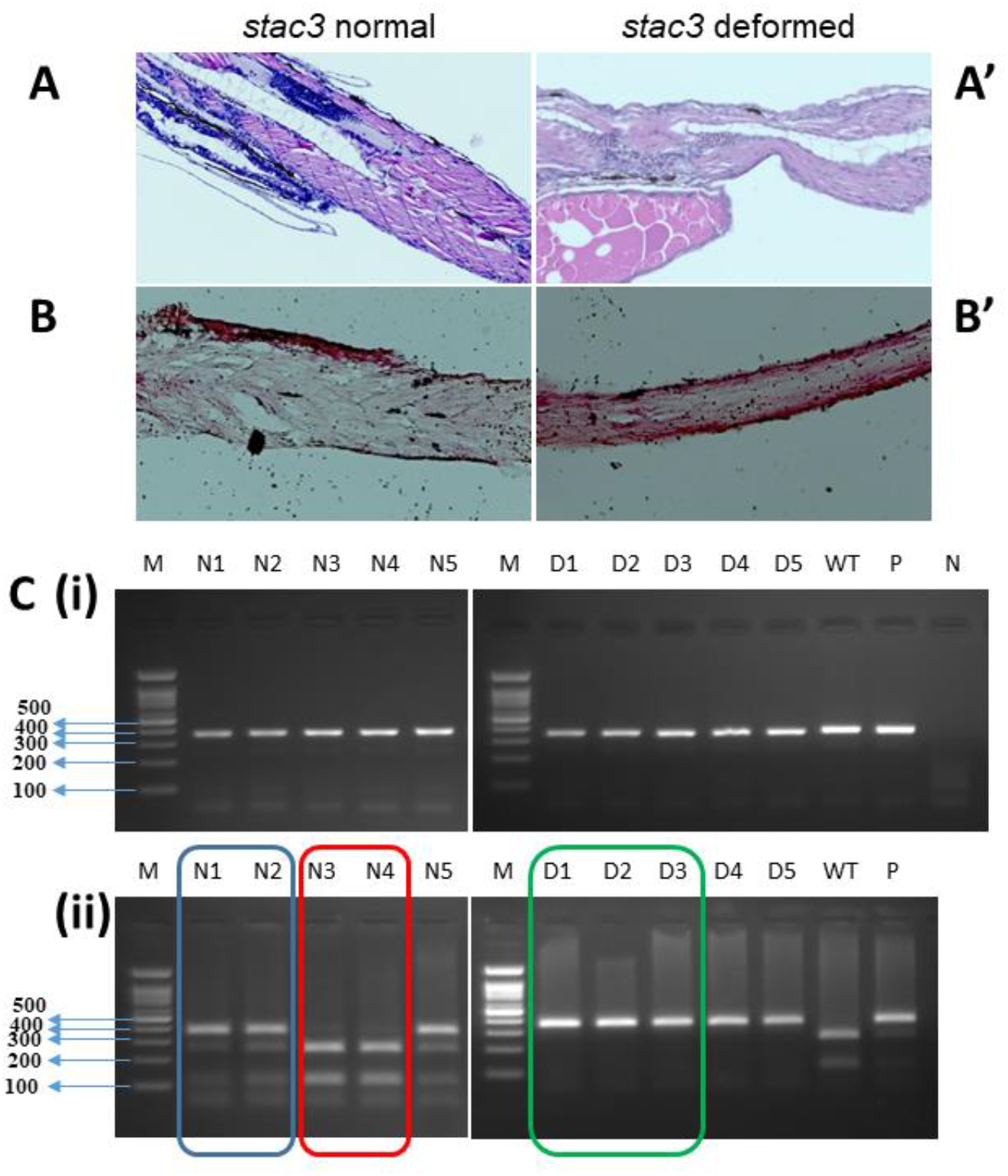
Muscle histology of *stac3*^-/-^ larvae. (A-A’). Paraffin based muscle sections of *stac3* wild type and heterozygous siblings and *stac3* knockout larvae stained with hematoxylin showed disorganized fibers in the knockout fish (A’). (B-B’). OCT based muscle sections of *stac3^+/+^* and *stac3^+/-^* control siblings and knockout larvae were stained with ORO dye. *stac3* knockout larvae displayed more red staining (higher lipids) compared to *stac3 stac3^+/+^* and *stac3^+/-^* control siblings at 5 dpf. (D). Genotype of larvae was confirmed as described in FigureS1A, B.

**S5.**
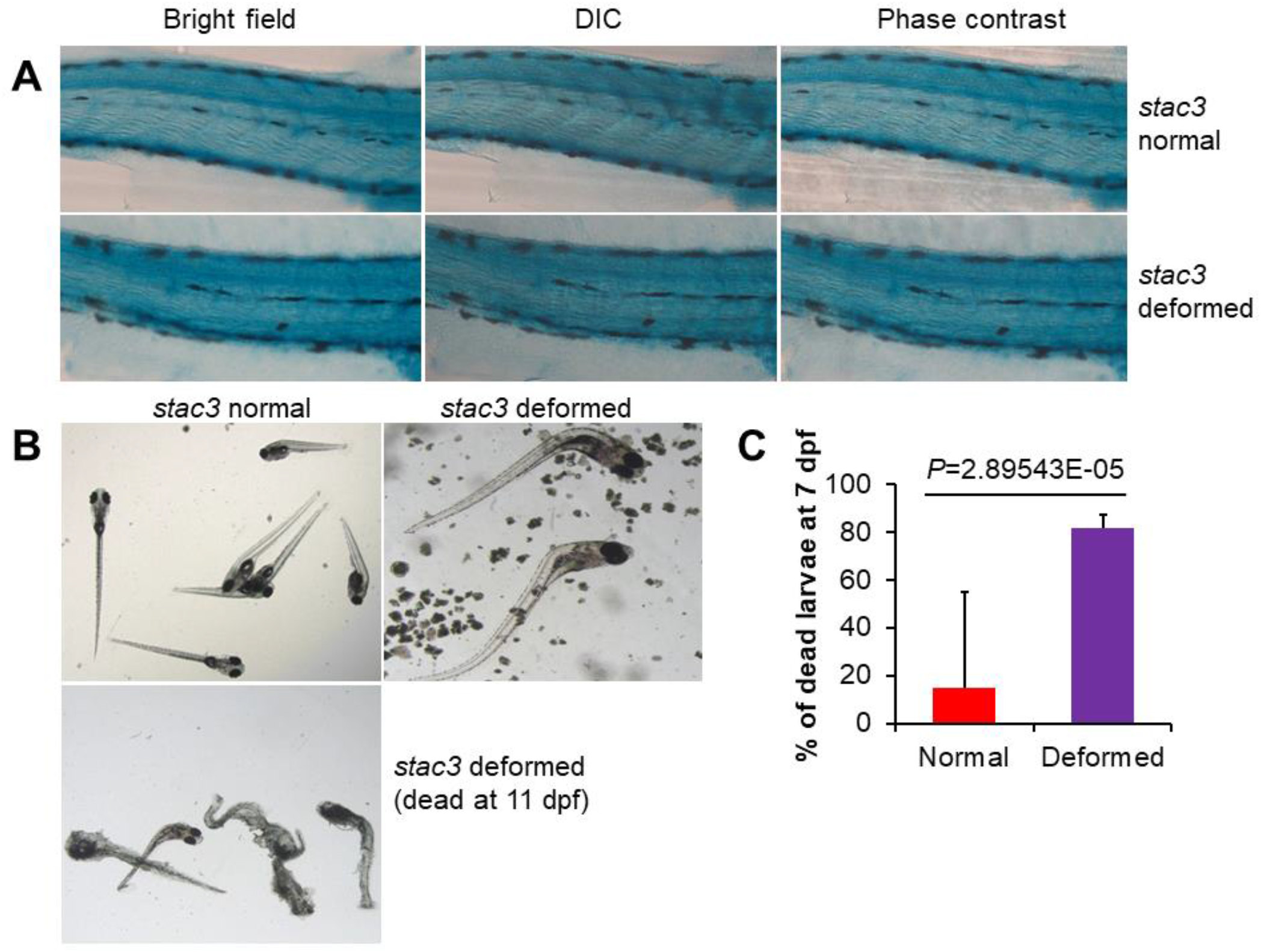
Death of *stac3^-/-^* larvae at the early age. (A). Altered integrity of muscle fibers observed in *stac3* KO larvae (bottom panel) compared to *stac3* wild type and heterozygous siblings (top panel), as measured by SA-β-gal staining at 6 dpf. (B). *stac3* wild type and heterozygous siblings and KO larvae raised in an incubator confirmed that stac3^-/-^ larvae expeditiously die by 11 dpf. stac3 wild type and heterozygous sibling (left: top) at 9 dpf, stac3 knockout (middle) 10 dpf, and stac3 knockout larvae (left: bottom) 11 dpf. (C). Percentage (%) calculation of dead *stac3* knockout larvae at 7 dpf. Scale bar: 0.5-1 mm.

**S6.**
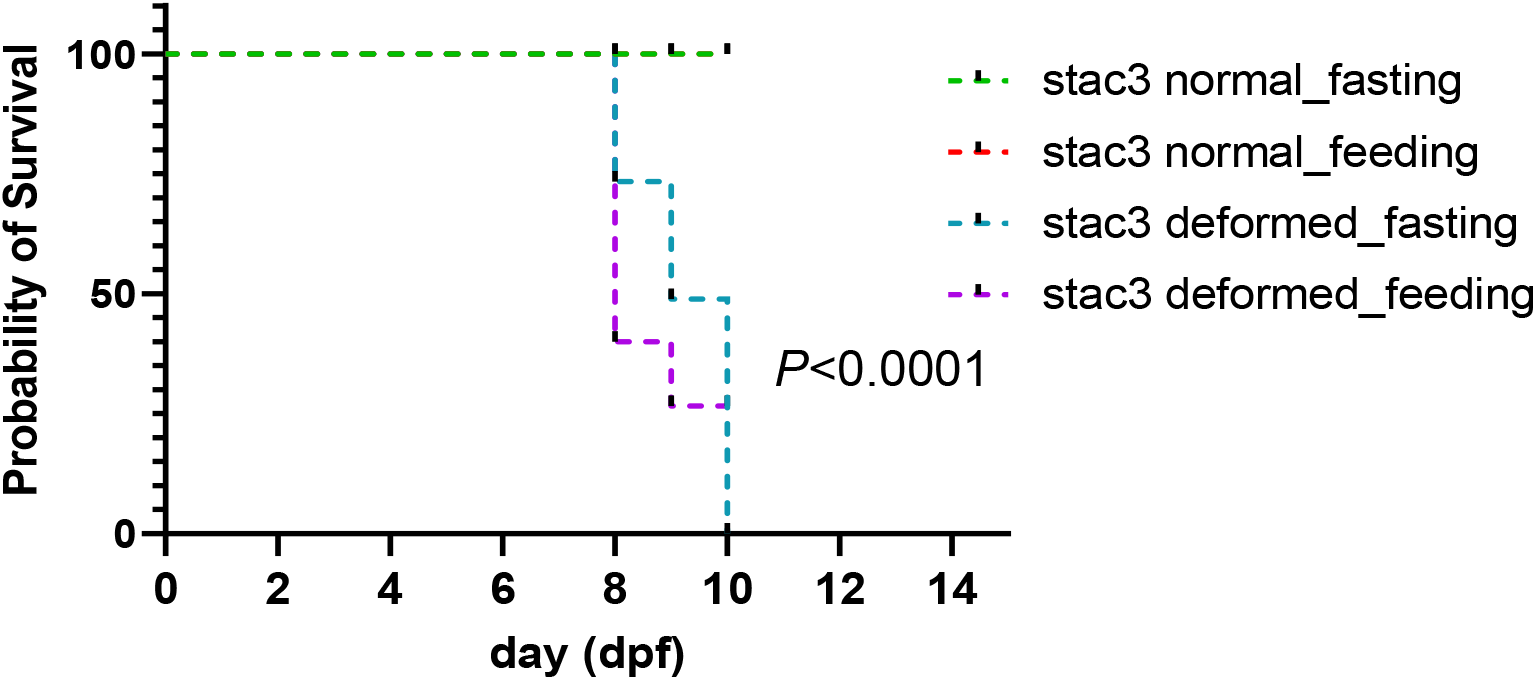
Fasted *stac3* KO larvae show a better short-term survival Feeding was introduced in some 7 dpf old *stac3* KO. (deformed) and *stac3*^+/+^ and *stac3*^+/-^ (normal) control siblings. Larval growth and survival were monitored for 72 hours and the fasted and fed animals were compared. Survival percentage was calculated between fasting vs. feed *stac3* KO, *stac3*^+/+^ and *stac3*^+/-^ groups using Log-rank test, ****p<0.0001.

**Supplementary table1.**
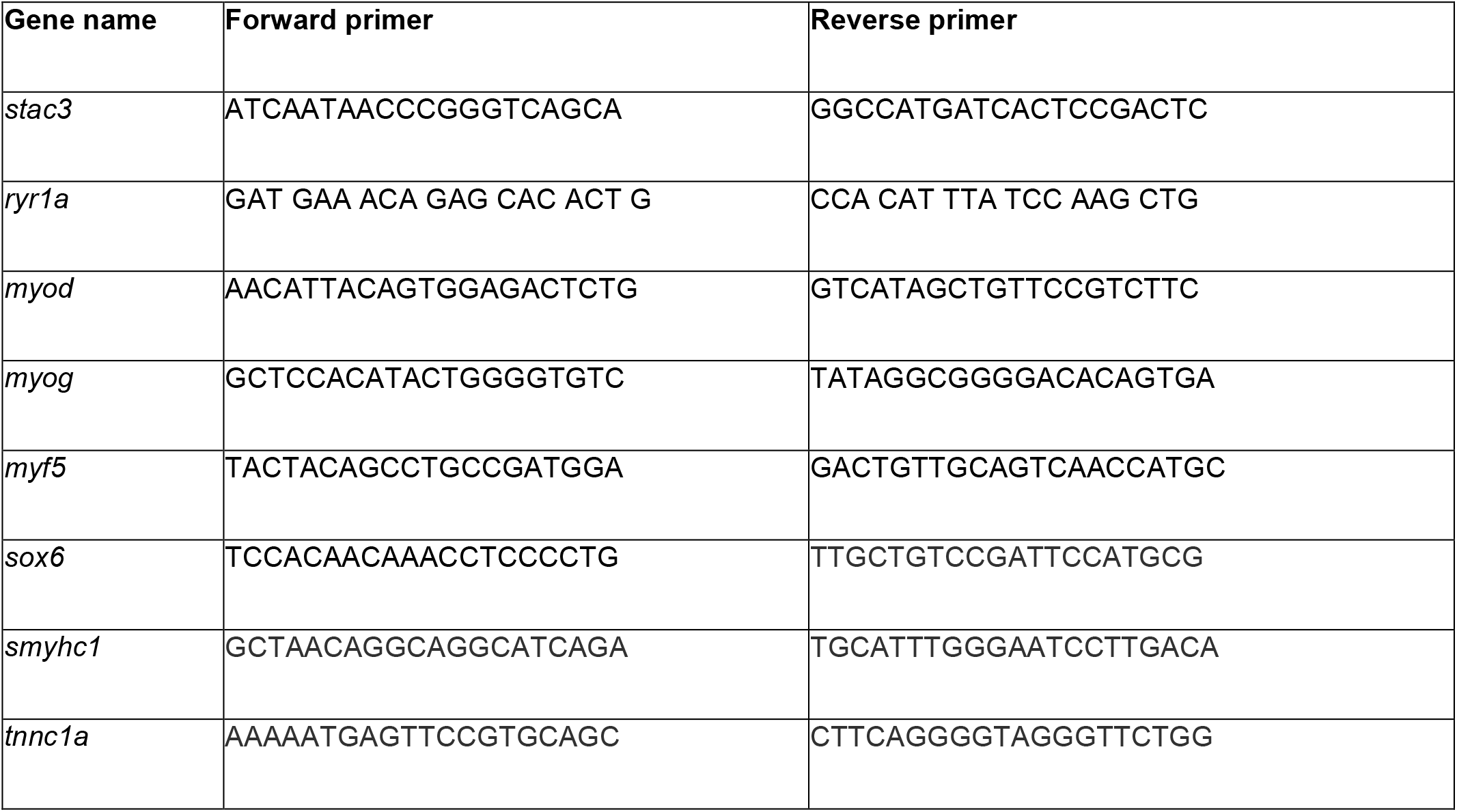
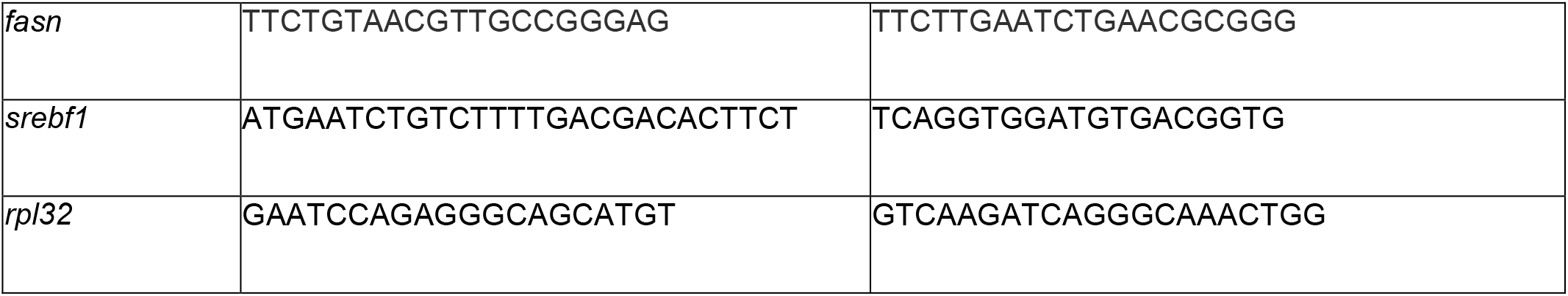
Primers used for RTqPCR.

**Supplementary table2.** Percentage (%) of dead *stac3* KO larvae at 7 dpf.

| Larva condition | Age: 7 dpf |  |  |  |  |  |
| --- | --- | --- | --- | --- | --- | --- |
|  | exp1 | exp2 | exp3 | Average | % in total population | % of dead |
| *stac3*<sup>+/+</sup> and *stac3*<sup>+/-</sup> control siblings | 324 | 379 | 300 | 334.33 | 87.14 | 14.76 |
| *stac3*<sup>-/-</sup> (KO) | 45 | 47 | 56 | 49.33 | 12.86 | 81.76 |
| *stac3* dead larvae (belong to KO group) | 45 | 47 | 29 | 40.33 | 10.51 |  |
| Total | 369 | 426 | 356 | 383.67 | 100.00 |  |

**Supplementary table3.**
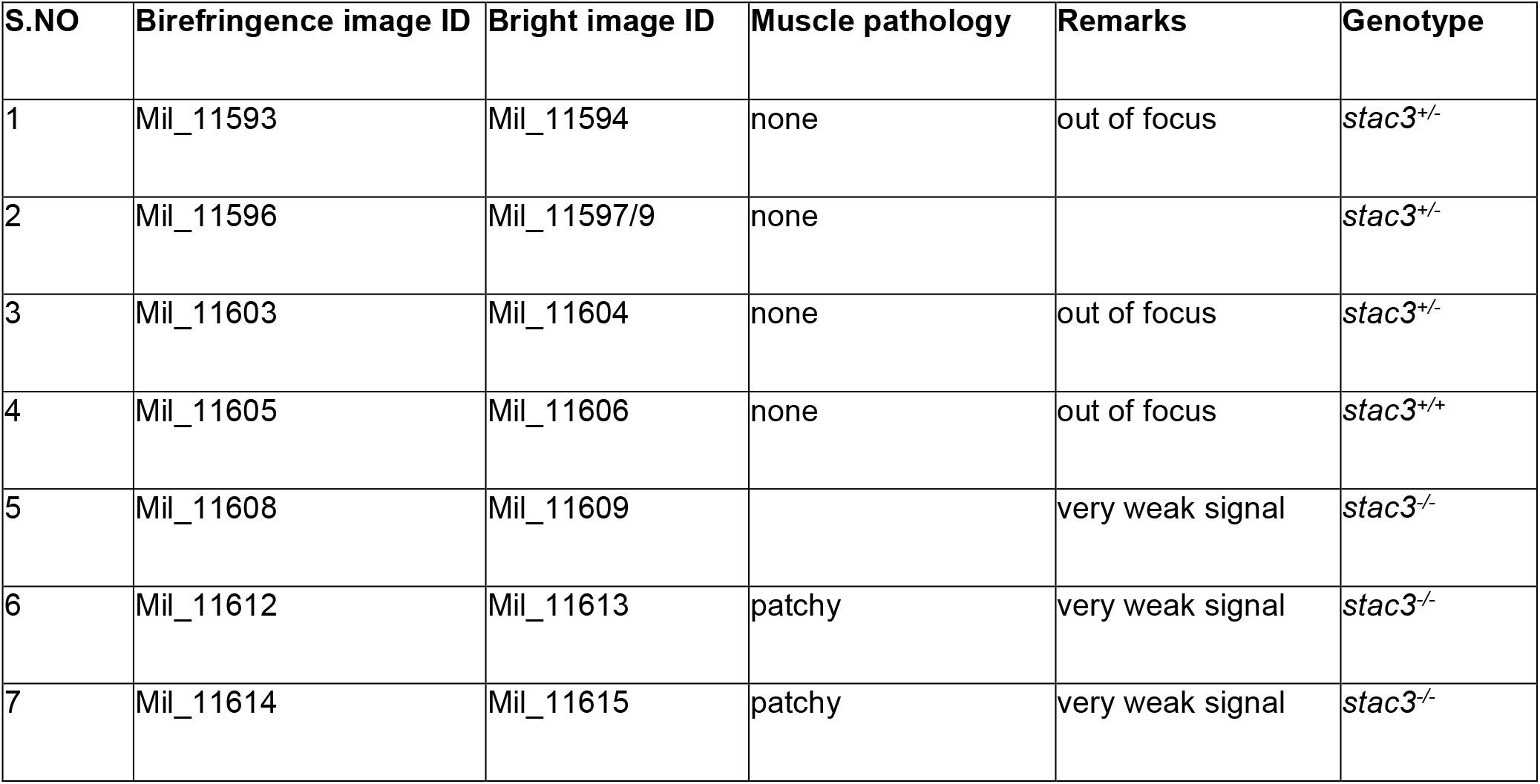
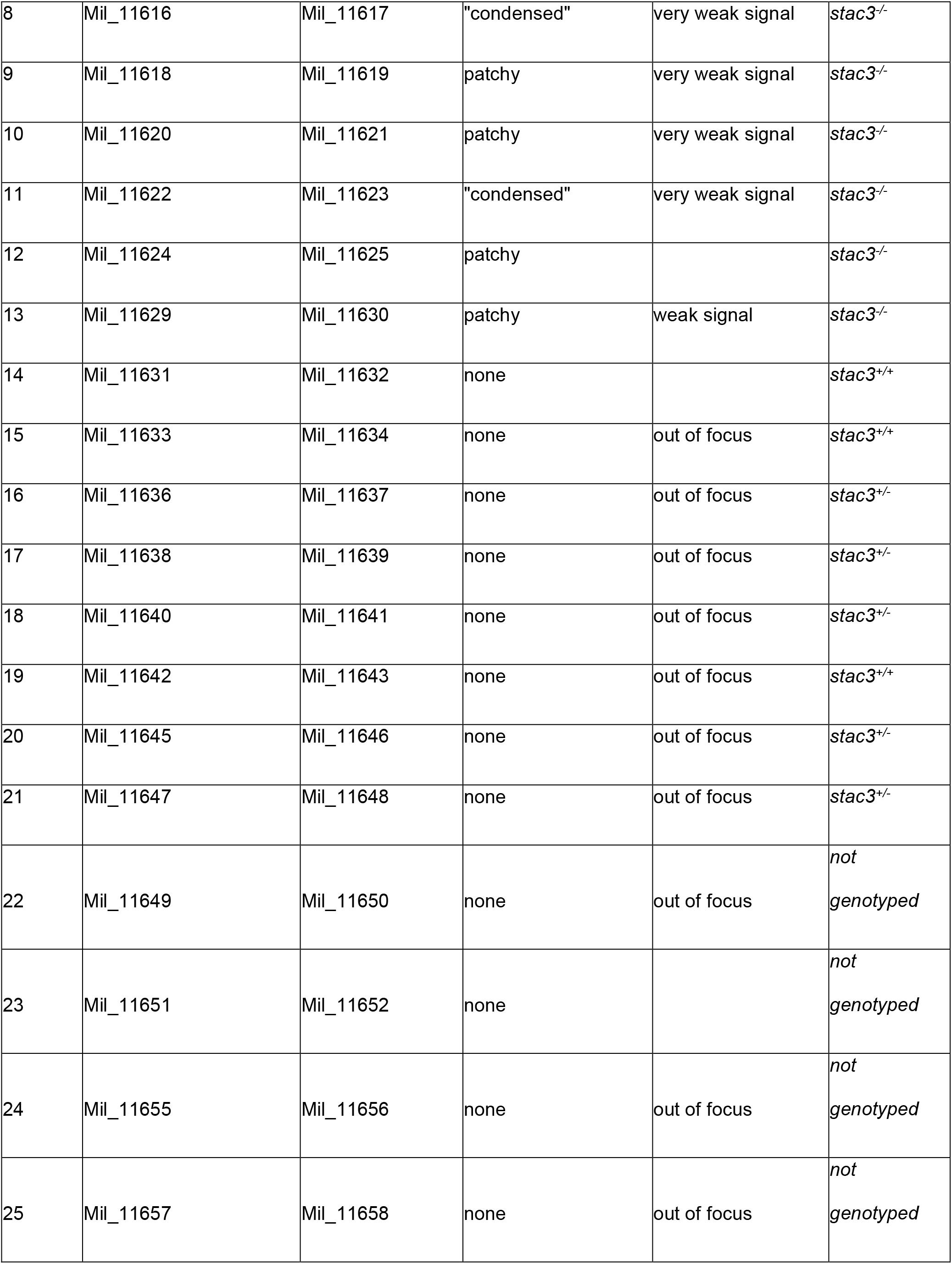
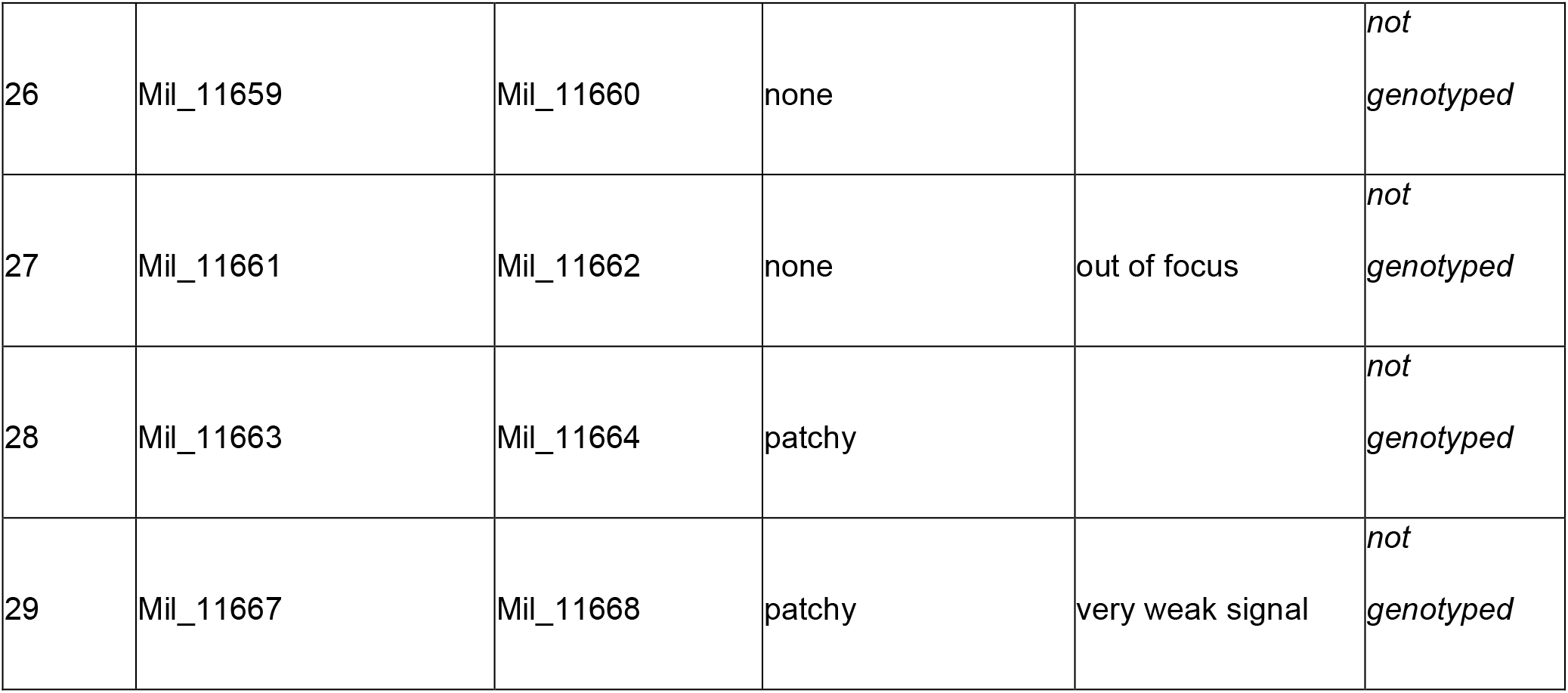
Muscle pathology identification by blind analysis of zebrafish larva at 6 dpf.

## References

1. McLeod, M., Breen, L., Hamilton, D.L., and Philp, A. Live strong and prosper: the importance of skeletal muscle strength for healthy ageing.

2. Gineste, C., and Laporte, J. (2023). Therapeutic approaches in different congenital myopathies. Current Opinion in Pharmacology 68, 102328.

3. Huang, K., Bi, F., and Huan, Y. (2021). A systematic review and meta-analysis of the prevalence of congenital myopathy. Frontiers in neurology, 1959.

4. Gineste, C., and Laporte, J. Therapeutic approaches in different congenital myopathies.

5. Tolchin, D., Yeager, J.P., Prasad, P., Dorrani, N., Russi, A.S., Martinez-Agosto, J.A., Haseeb, A., Angelozzi, M., Santen, G.W.E., Ruivenkamp, C., et al. De Novo SOX6 Variants Cause a Neurodevelopmental Syndrome Associated with ADHD, Craniosynostosis, and Osteochondromas.

6. Claeys, K.G. (2020). Congenital myopathies: an update. Developmental Medicine & Child Neurology 62, 297–302.

7. Ogasawara, M., and Nishino, I. (2023). A review of major causative genes in congenital myopathies. Journal of Human Genetics 68, 215–225.

8. Mendieta-Serrano, M.A., Dhar, S., Ng, B.H., Narayanan, R., Lee, J.J.Y., Ong, H.T., Toh, P.J.Y., Röllin, A., Roy, S., and Saunders, T.E. (2022). Slow muscles guide fast myocyte fusion to ensure robust myotome formation despite the high spatiotemporal stochasticity of fusion events. Developmental Cell 57, 2095–2110.e2095.

9. Zempo, B., Yamamoto, Y., Williams, T., and Ono, F. Synaptic silencing of fast muscle is compensated by rewired innervation of slow muscle. Science Advances 6, eaax8382.

10. Jackson, H.E., Ono, Y., Wang, X., Elworthy, S., Cunliffe, V.T., and Ingham, P.W. (2015). The role of Sox6 in zebrafish muscle fiber type specification. Skeletal Muscle 5, 2.

11. Li, S., Wen, H., and Du, S. Defective sarcomere organization and reduced larval locomotion and fish survival in slow muscle heavy chain 1 (smyhc1) mutants.

12. Stamm, D.S., Aylsworth, A.S., Stajich, J.M., Kahler, S.G., Thorne, L.B., Speer, M.C., and Powell, C.M. (2008). Native American myopathy: Congenital myopathy with cleft palate, skeletal anomalies, and susceptibility to malignant hyperthermia. American Journal of Medical Genetics Part A 146A, 1832-1841.

13. Miller, D.M., Daly, C., Aboelsaod, E.M., Gardner, L., Hobson, S.J., Riasat, K., Shepherd, S., Robinson, R.L., Bilmen, J.G., Gupta, P.K., et al. Genetic epidemiology of malignant hyperthermia in the UK.

14. Rufenach, B., and Van Petegem, F. Structure and function of STAC proteins: Calcium channel modulators and critical components of muscle excitation-contraction coupling.

15. Cong, X., Doering, J., Mazala, D.A., Chin, E.R., Grange, R.W., and Jiang, H. (2016). The SH3 and cysteine-rich domain 3 (Stac3) gene is important to growth, fiber composition, and calcium release from the sarcoplasmic reticulum in postnatal skeletal muscle. Skeletal muscle 6, 1–17.

16. Bower, N.I., de la Serrana Dg Fau - Cole, N.J., Cole Nj Fau - Hollway, G.E., Hollway Ge Fau - Lee, H.-T., Lee Ht Fau - Assinder, S., Assinder S Fau - Johnston, I.A., and Johnston, I.A. Stac3 is required for myotube formation and myogenic differentiation in vertebrate skeletal muscle.

17. Saini-Chohan, H.K., Mitchell Rw Fau - Vaz, F.M., Vaz Fm Fau - Zelinski, T., Zelinski T Fau - Hatch, G.M., and Hatch, G.M. Delineating the role of alterations in lipid metabolism to the pathogenesis of inherited skeletal and cardiac muscle disorders: Thematic Review Series: Genetics of Human Lipid Diseases.

18. Seyssel, K., Alligier, M., Meugnier, E., Chanseaume, E., Loizon, E., Canto, C., Disse, E., Lambert-Porcheron, S., Brozek, J., Blond, E., et al. (2014). Regulation of Energy Metabolism and Mitochondrial Function in Skeletal Muscle During Lipid Overfeeding in Healthy Men. The Journal of Clinical Endocrinology & Metabolism 99, E1254–E1262.

19. Shimano, H., and Sato, R. (2017). SREBP-regulated lipid metabolism: convergent physiology — divergent pathophysiology. Nature Reviews Endocrinology 13, 710–730.

20. Linsley, J.W., Hsu, I.-U., Groom, L., Yarotskyy, V., Lavorato, M., Horstick, E.J., Linsley, D., Wang, W., Franzini-Armstrong, C., and Dirksen, R.T. (2017). Congenital myopathy results from misregulation of a muscle Ca2+ channel by mutant Stac3. Proceedings of the National Academy of Sciences 114, E228–E236.

21. Rufenach, B., Christy, D., Flucher, B.E., Bui, J.M., Gsponer, J., Campiglio, M., and Van Petegem, F. Multiple Sequence Variants in STAC3 Affect Interactions with Ca(V)1.1 and Excitation-Contraction Coupling.

22. Horstick, E.J., Linsley, J.W., Dowling, J.J., Hauser, M.A., McDonald, K.K., Ashley-Koch, A., Saint-Amant, L., Satish, A., Cui, W.W., and Zhou, W. (2013). Stac3 is a component of the excitation–contraction coupling machinery and mutated in Native American myopathy. Nature communications 4, 1952.

23. Berger, J., and Currie, P.D. Zebrafish models flex their muscles to shed light on muscular dystrophies.

24. Sztal, T.E., Zhao, M., Williams, C., Oorschot, V., Parslow, A.C., Giousoh, A., Yuen, M., Hall, T.E., Costin, A., Ramm, G., et al. (2015). Zebrafish models for nemaline myopathy reveal a spectrum of nemaline bodies contributing to reduced muscle function. Acta Neuropathologica 130, 389–406.

25. Daya, A., Donaka, R., and Karasik, D. (2020). Zebrafish models of sarcopenia. Disease Models & Mechanisms 13, dmm042689.

26. Mesika, A., Nadav, G., Shochat, C., Kalfon, L., Jackson, K., Khalaileh, A., Karasik, D., and Falik-Zaccai, T.C. (2022). NGLY1 Deficiency Zebrafish Model Manifests Abnormalities of the Nervous and Musculoskeletal Systems. Frontiers in Cell and Developmental Biology 10.

27. Li, H., Xu, J., Bian, Y.-H., Rotllant, P., Shen, T., Chu, W., Zhang, J., Schneider, M., and Du, S.J. (2011). Smyd1b_tv1, a Key Regulator of Sarcomere Assembly, Is Localized on the M- Line of Skeletal Muscle Fibers. PLOS ONE 6, e28524.

28. Kishi, S., Bayliss, P.E., Uchiyama, J., Koshimizu, E., Qi, J., Nanjappa, P., Imamura, S., Islam, Neuberg, D., Amsterdam, A., et al. (2008). The Identification of Zebrafish Mutants Showing Alterations in Senescence-Associated Biomarkers. PLOS Genetics 4, e1000152.

29. Livne, H., Avital, T., Ruppo, S., Harazi, A., Mitrani-Rosenbaum, S., and Daya, A. (2022).Generation and characterization of a novel gne Knockout Model in Zebrafish. Frontiers in Cell and Developmental Biology 10.

30. Sokoloff, A.J., Yang, B., Li, H., and Burkholder, T.J. (2007). Immunohistochemical characterization of slow and fast myosin heavy chain composition of muscle fibres in the styloglossus muscle of the human and macaque (Macaca rhesus). Arch Oral Biol 52, 533–543.

31. Stewart, M.A., Franks-Skiba, K., Chen, S., and Cooke, R. (2010). Myosin ATP turnover rate is a mechanism involved in thermogenesis in resting skeletal muscle fibers. Proc Natl Acad Sci U S A 107, 430–435.

32. Yamamoto, M., Legendre, N.P., Biswas, A.A., Lawton, A., Yamamoto, S., Tajbakhsh, S., Kardon, G., and Goldhamer, D.J. Loss of MyoD and Myf5 in Skeletal Muscle Stem Cells Results in Altered Myogenic Programming and Failed Regeneration.

33. Cao, Y., Kumar Rm Fau - Penn, B.H., Penn Bh Fau - Berkes, C.A., Berkes Ca Fau - Kooperberg, C., Kooperberg C Fau - Boyer, L.A., Boyer La Fau - Young, R.A., Young Ra Fau - Tapscott, S.J., and Tapscott, S.J. Global and gene-specific analyses show distinct roles for Myod and Myog at a common set of promoters.

34. Reinholt, B.M., Ge, X., Cong, X., Gerrard, D.E., and Jiang, H. (2013). Stac3 is a novel regulator of skeletal muscle development in mice. PloS one 8, e62760.

35. Shi, H., Verma, M., Zhang, L., Dong, C., Flavell, R.A., and Bennett, A.M. (2013). Improved regenerative myogenesis and muscular dystrophy in mice lacking Mkp5. The Journal of Clinical Investigation 123, 2064–2077.

36. Quinlivan, V.H., and Farber, S.A. (2017). Lipid Uptake, Metabolism, and Transport in the Larval Zebrafish. Frontiers in Endocrinology 8.

37. Jiang, K., Pichol-Thievend, C., Neufeld, Z., and Francois, M. Assessment of heterogeneity in collective endothelial cell behavior with multicolor clonal cell tracking to predict arteriovenous remodeling.

38. Stamm, D.S., Aylsworth As Fau - Stajich, J.M., Stajich Jm Fau - Kahler, S.G., Kahler Sg Fau - Thorne, L.B., Thorne Lb Fau - Speer, M.C., Speer Mc Fau - Powell, C.M., and Powell, C.M. Native American myopathy: congenital myopathy with cleft palate, skeletal anomalies, and susceptibility to malignant hyperthermia.

39. Kepser, L.-J., Damar, F., De Cicco, T., Chaponnier, C., Prószyński, T.J., Pagenstecher, A., and Rust, M.B. (2019). CAP2 deficiency delays myofibril actin cytoskeleton differentiation and disturbs skeletal muscle architecture and function. Proceedings of the National Academy of Sciences 116, 8397–8402.

40. Barcelos, R.P., Lima, F.D., Carvalho, N.R., Bresciani, G., and Royes, L.F. Caffeine effects on systemic metabolism, oxidative-inflammatory pathways, and exercise performance.

41. Tallis, J., James Rs Fau - Cox, V.M., Cox Vm Fau - Duncan, M.J., and Duncan, M.J. The effect of physiological concentrations of caffeine on the power output of maximally and submaximally stimulated mouse EDL (fast) and soleus (slow) muscle.

42. Gonçalves, D.F., Tassi, C.C., Amaral, G.P., Stefanello, S.T., Dalla Corte, C.L., Soares, F.A., Posser, T., Franco, J.L., and Carvalho, N.R. Effects of caffeine on brain antioxidant status and mitochondrial respiration in acetaminophen-intoxicated mice.

43. Nelson, B.R., Wu, F., Liu, Y., Anderson, D.M., McAnally, J., Lin, W., Cannon, S.C., Bassel- Duby, R., and Olson, E.N. (2013). Skeletal muscle-specific T-tubule protein STAC3 mediates voltage-induced Ca2+ release and contractility. Proceedings of the National Academy of Sciences 110, 11881–11886.

44. Conerly, M.L., Yao, Z., Zhong, J.W., Groudine, M., and Tapscott, S.J. Distinct Activities of Myf5 and MyoD Indicate Separate Roles in Skeletal Muscle Lineage Specification and Differentiation.

45. Ganassi, M., Badodi, S., Wanders, K., Zammit, P.S., and Hughes, S.M. (2020). Myogenin is an essential regulator of adult myofibre growth and muscle stem cell homeostasis. eLife 9, e60445.

46. Mok, G.F., Mohammed, R.H., and Sweetman, D. Expression of myogenic regulatory factors in chicken embryos during somite and limb development.

47. Malila, Y., Thanatsang, K.V., Sanpinit, P., Arayamethakorn, S., Soglia, F., Zappaterra, M., Bordini, M., Sirri, F., Rungrassamee, W., Davoli, R., et al. (2022). Differential expression patterns of genes associated with metabolisms, muscle growth and repair in Pectoralis major muscles of fast- and medium-growing chickens. PLOS ONE 17, e0275160.

48. Wen, C., Jiang, X., Ding, L., Wang, T., and Zhou, Y. (2017). Effects of dietary methionine on breast muscle growth, myogenic gene expression and IGF-I signaling in fast- and slow- growing broilers. Scientific Reports 7, 1924.

49. Coppens, S., Barnard, A.M., Puusepp, S., Pajusalu, S., Õunap, K., Vargas-Franco, D., Bruels, C.C., Donkervoort, S., Pais, L., Chao, K.R., et al. (2021). A form of muscular dystrophy associated with pathogenic variants in JAG2. The American Journal of Human Genetics 108, 840–856.

50. Jungbluth, H., Ochala, J., Treves, S., and Gautel, M. (2017). Current and future therapeutic approaches to the congenital myopathies. Seminars in Cell & Developmental Biology 64, 191–200.

51. Schessl, J., Taratuto, A.L., Sewry, C., Battini, R., Chin, S.S., Maiti, B., Dubrovsky, A.L., Erro, M.G., Espada, G., Robertella, M., et al. (2009). Clinical, histological and genetic characterization of reducing body myopathy caused by mutations in FHL1. Brain 132, 452–464.

52. Böhm, J., Vasli, N., Malfatti, E., Le Gras, S., Feger, C., Jost, B., Monnier, N., Brocard, J., Karasoy, H., Gérard, M., et al. (2013). An Integrated Diagnosis Strategy for Congenital Myopathies. PLOS ONE 8, e67527.

53. Cassandrini, D., Trovato, R., Rubegni, A., Lenzi, S., Fiorillo, C., Baldacci, J., Minetti, C., Astrea, G., Bruno, C., Santorelli, F.M., et al. (2017). Congenital myopathies: clinical phenotypes and new diagnostic tools. Italian Journal of Pediatrics 43, 101.

54. Stamm, D., Powell, C., Stajich, J., Zismann, V., Stephan, D., Chesnut, B., Aylsworth, A., Kahler, S., Deak, K., and Gilbert, J. (2008). Novel congenital myopathy locus identified in Native American Indians at 12q 13. 13-14.1. Neurology 71, 1764-1769.

55. Talbot, J., and Maves, L. Skeletal muscle fiber type: using insights from muscle developmental biology to dissect targets for susceptibility and resistance to muscle disease.

56. Miyares, R.L., de Rezende, V.B., and Farber, S.A. Zebrafish yolk lipid processing: a tractable tool for the study of vertebrate lipid transport and metabolism.

57. Fraher, D., Sanigorski, A., Mellett, Natalie A., Meikle, Peter J., Sinclair, Andrew J., and Gibert, Y. (2016). Zebrafish Embryonic Lipidomic Analysis Reveals that the Yolk Cell Is Metabolically Active in Processing Lipid. Cell Reports 14, 1317–1329.

58. Goi, M., and Childs, S.J. (2016). Patterning mechanisms of the sub-intestinal venous plexus in zebrafish. Developmental Biology 409, 114–128.

59. Edinburgh, R.M., Bradley, H.E., Abdullah, N.-F., Robinson, S.L., Chrzanowski-Smith, O.J., Walhin, J.-P., Joanisse, S., Manolopoulos, K.N., Philp, A., Hengist, A., et al. (2020). Lipid Metabolism Links Nutrient-Exercise Timing to Insulin Sensitivity in Men Classified as Overweight or Obese. The Journal of Clinical Endocrinology & Metabolism 105, 660–676.

60. Goodpaster, B.H., Bergman, B.C., Brennan, A.M., and Sparks, L.M. (2022). Intermuscular adipose tissue in metabolic disease. Nature Reviews Endocrinology.

61. Yoganantharjah, P., Byreddy Ar Fau - Fraher, D., Fraher D Fau - Puri, M., Puri M Fau - Gibert, Y., and Gibert, Y. Rapid quantification of neutral lipids and triglycerides during zebrafish embryogenesis.

62. Sinha, S., Elbaz-Alon, Y., and Avinoam, O. (2022). Ca2+ as a coordinator of skeletal muscle differentiation, fusion and contraction. The FEBS Journal 289, 6531–6542.

63. Telegrafi, A, Webb, B, Robbins, S.M., Speck-Martins, C.E., FitzPatrick, D., Fleming, L.A, Redett, R., Dufke, A., Houge, G, van Harssel, J.J.T., et al. Identification of STAC3 variants in non-Native American families with overlapping features of Carey-Fineman-Ziter syndrome and Moebius syndrome.

64. Gromand, M., Gueguen, P., Pervillé, A., Ferroul, F., Morel, G., Harouna, A., Doray, B., Urtizberea, J.A., Alessandri, J.-L., and Robin, S. (2022). STAC3 related congenital myopathy: A case series of seven Comorian patients. European Journal of Medical Genetics 65, 104598.

65. Murtazina, A., Demina, N., Chausova, P., Shchagina, O., Borovikov, A., and Dadali, E. (2022). The First Russian Patient with Native American Myopathy. In Genes.

66. Patton, E.E., Zon, L.I., and Langenau, D.M. (2021). Zebrafish disease models in drug discovery: from preclinical modelling to clinical trials. Nature Reviews Drug Discovery 20, 611–628.

67. Espinosa, K.G., Geissah, S., Groom, L., Volpatti, J., Scott, I, Dirksen, R.T., Zhao, M, and Dowling, J. Characterization of a novel zebrafish model of SPEG-related centronuclear myopathy. LID - 10.1242/dmm.049437.

